# PPM1M, a LRRK2-counteracting, phosphoRab12-preferring phosphatase with potential link to Parkinson’s disease

**DOI:** 10.1101/2025.03.19.644182

**Authors:** Claire Y. Chiang, Neringa Pratuseviciute, Yu-En Lin, Ayan Adhikari, Wondwossen M. Yeshaw, Chloe Flitton, Pemba L. Sherpa, Francesca Tonelli, Irena Rektorova, Timothy Lynch, Joanna Siuda, Monika Rudzińska-Bar, Oleksandr Pulyk, Peter Bauer, Christian Beetz, Dennis W. Dickson, Owen A. Ross, Zbigniew K. Wszolek, Christine Klein, Alexander Zimprich, Dario R. Alessi, Esther M. Sammler, Suzanne R. Pfeffer

## Abstract

Leucine-rich repeat kinase 2 (LRRK2) phosphorylates a subset of Rab GTPases that regulate receptor trafficking; activating mutations in *LRRK2* are linked to Parkinson’s disease. Rab phosphorylation is a transient event that can be reversed by phosphatases, including PPM1H, that acts on phosphoRab8A and phosphoRab10. Here we report a phosphatome-wide siRNA screen that identified PPM1M as a phosphoRab12-preferring phosphatase that also acts on phosphoRab8A and phosphoRab10. Upon knockout from cells or mice, PPM1M displays selectivity for phosphoRab12. As shown previously for mice harboring LRRK2 pathway mutations, knockout of *Ppm1m* leads to primary cilia loss in striatal cholinergic interneurons. We have also identified a rare *PPM1M* mutation in patients with Parkinson’s disease that is catalytically inactive when tested *in vitro* and in cells. These findings identify PPM1M as a key player in the LRRK2 signaling pathway and provide a new therapeutic target for the possible benefit of patients with Parkinson’s disease.

**Teaser:** Parkinson’s linked Rab phosphorylation is reversed by PPM1M; the inactive D440N variant is implicated in rare patient cases.

## Introduction

While the cause of most Parkinson’s disease (PD) cases is unknown, perhaps as many as 20% have a strong genetic basis (*1, 2*). Of these, around 3% are due to activating mutations in the *LRRK2* gene (*3*), making it one of the most clinically relevant PD genes. *LRRK2* pathogenic mutations display population-specific frequencies (*2, 4*). The most common global pathogenic *LRRK2* mutation is G2019S, which increases kinase activity about two-fold (*5*). *LRRK2* G2019S has a high frequency in Ashkenazi Jewish (15-20%) and North African Berber-Arab (>30%) patients with PD (*2*). Other population-specific variants include *LRRK2* R1441G (Basque population, ∼6%) and R1628P and G2385R (East Asian population, 5-10% of patients; two-fold risk increase) (*2, 4*). Notably, a protective *LRRK2* haplotype (N551K-R1398H-K1423K) has also been identified, suggesting kinase activity—and therefore disease risk—is modifiable (*4, 6*). LRRK2 is considered a highly promising target for PD treatment, and several clinical trials are underway (*7*). Unlike current PD treatments that primarily address symptoms, LRRK2-specific kinase inhibitors could potentially slow or halt PD progression. Moreover, LRRK2 inhibitors would likely benefit patients with both *LRRK2* mutant and idiopathic forms of PD.

LRRK2 phosphorylates a subset of the 65 human Rab GTPases (*8, 9*) that are master regulators of membrane trafficking (*10, 11*). Upon phosphorylation, these Rab GTPases bind to a new set of phosphoRab-specific effector proteins to drive cellular pathology (*9*). In 2019, Berndsen et al. (*12*) reported the discovery of PPM1H (Protein Phosphatase, Mg2^+^/Mn2^+^ Dependent 1H) as a Rab GTPase-specific, LRRK2 action-reversing phosphatase. PPM1H shows a preference for Rab8A, Rab10, and Rab35, and dephosphorylation of these Rabs regulates primary cilia formation (*12*) and autophagy (*13*). PPM1H relies on an N-terminal amphipathic helix for Golgi localization in cells (*14*) and under certain conditions it can be detected on mitochondrial surfaces (*15*). PPM1H is most active when associated with highly curved membranes, where interaction of its N-terminal amphipathic helix with those membranes enhances its catalytic activity (*14*). PPM1H’s specificity for LRRK2-phosphorylated Rab proteins is largely determined by a unique structural feature called a “flap” domain (*16*). Domain swap experiments have shown that this 110-residue domain, located adjacent to the active site, has evolved to recognize specific phosphorylated Rab GTPases (hereafter referred to as phosphoRabs) (*16*). PPM1H’s ability to reverse pathological Rab phosphorylation positions it as another potential therapeutic target for PD, with activity enhancers offering a novel approach to modulate LRRK2 signaling and mitigate disease progression.

While PPM1H is highly specific for certain Rab proteins, it does not act efficiently on all LRRK2-phosphorylated Rabs. For example, Rab12 is a poor substrate for PPM1H in cellular contexts (*12*), suggesting that other phosphatases may also play a role. Thus, we carried out two additional, siRNA-based, phosphatome-wide screens to identify additional Rab-specific phosphatases. We present here the discovery of the PPM1H-related PPM1M as a phosphoRab12-preferring, Rab-specific phosphatase that has a rare, PD-associated, catalytically inactivating mutation.

## Results

### PPM1H does not act alone: evidence for additional Rab phosphatases

Detailed analysis revealed that PPM1H activity is not sufficient to explain phosphoRab10 dephosphorylation kinetics. To measure PPM1H-dependent dephosphorylation, we compared phosphoRab levels in cells with and without PPM1H protein; the LRRK2 kinase inhibitor, MLi-2 (*17*), was added for various times to block Rab protein re-phosphorylation (Fig. 1A). At baseline, phosphoRab10 levels were three-fold higher in PPM1H knockout cells relative to wild-type (WT) cells (Fig. 1B, red versus gray symbols), while phosphoRab12 levels were unchanged (Fig. 1B, blue and black symbols). Thus, as we have reported previously (*12*), phosphoRab12 levels are not impacted by loss of PPM1H.

**Figure 1.**
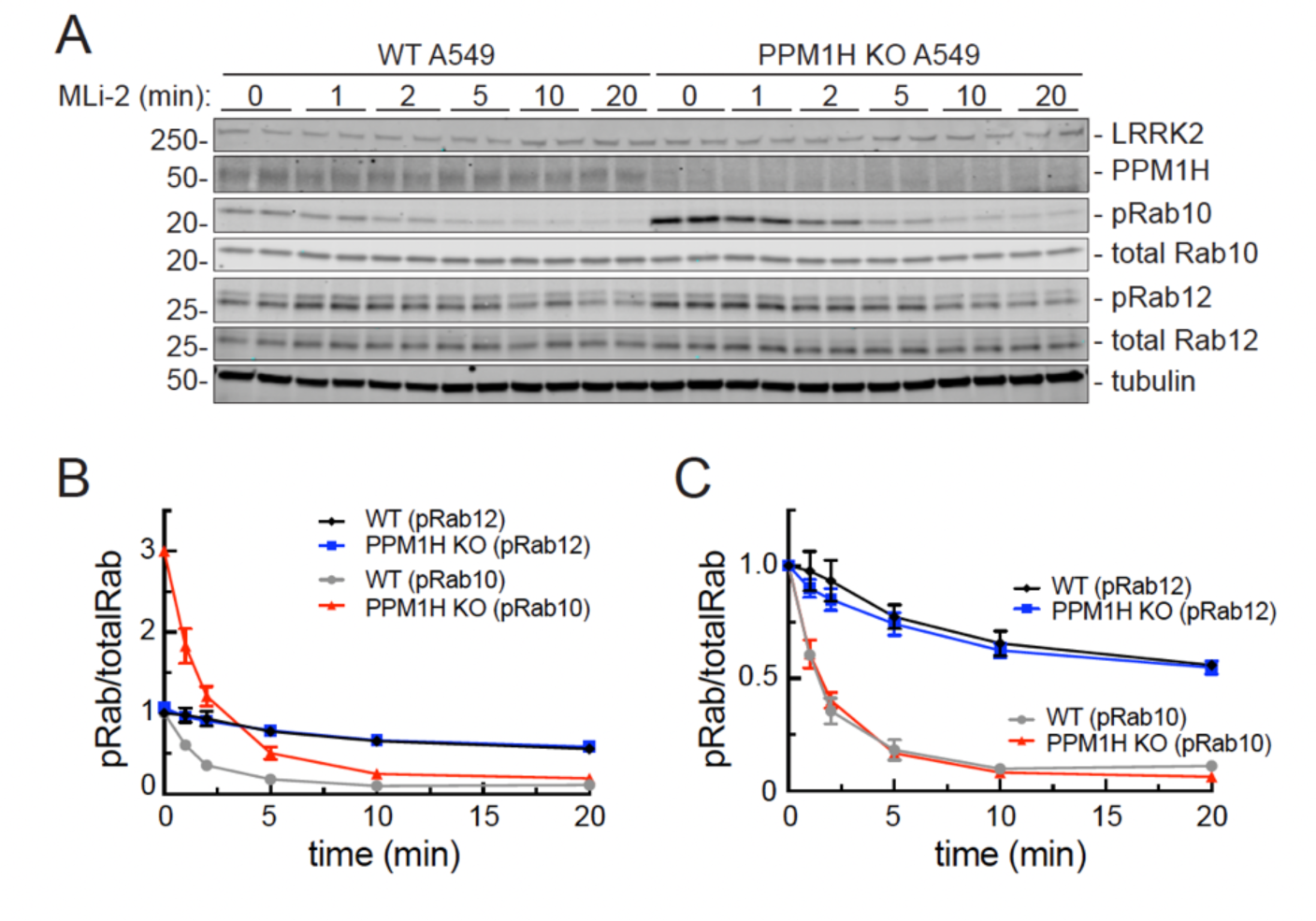
PPM1H knockout does not influence the rate of phosphoRab turnover. **(A)** Immunoblot analysis of wild-type (WT) and PPM1H knockout A549 cells, treated with 100 nM MLi-2 for the times indicated. Mass is shown at left in kDa here and in all subsequent figures; antigens are indicated at right. **(B)** Quantitation of pRab10/total Rab10 and pRab12/total Rab12 levels from immunoblots in (A), normalized to 1.0 for respective pRab10 or pRab12 WT 0 min conditions, as indicated. **(C)** Quantitation of pRab10/total Rab10 and pRab12/total Rab12 levels from immunoblots in (A), normalized to 1.0 for 0 min of each respective (WT or KO) condition to permit direct comparison. Error bars represent SEM from 3 independent experiments carried out in duplicate.

Normalization of the data for direct comparison revealed that both phosphoRab10 and phosphoRab12 continue to be dephosphorylated at equal rates, irrespective of the presence of PPM1H (Fig. 1C). Under these conditions, phosphoRab10 turned over very quickly, with a half-life of about two minutes, while phosphoRab12 displayed a half-life of 20 minutes. These data demonstrate the existence of another phosphoRab10 phosphatase(s), at least in A549 cells, with preference for phosphoRab10 over phosphoRab12.

### Phosphatome-wide screen identifies PPM1M as another phosphoRab phosphatase

We performed a phosphatome-wide siRNA screen to identify a novel phosphoRab12 phosphatase and/or a second phosphoRab10 phosphatase. Our screen compared 303 siRNAs targeting phosphatases and corresponding regulatory subunits in mouse 3T3 cells (Fig. 2A, Supp. Table 2). 3T3 cells were selected because they have performed well in prior CRISPR screens (*18*) and, importantly, they contain sufficient LRRK2 to enable detection of endogenous phosphoRab10 and phosphoRab12. To identify a LRRK2-counteracting phosphoRab12 phosphatase, we chose a longer duration of MLi-2 treatment (20 min) than used previously (5 min for phosphoRab10 in Berndsen et al. (*12*)) to best evaluate phosphoRab12 turnover, as phosphoRab12 turns over more slowly than phosphoRab10 (Fig. 1). After 72 hours with siRNA, cells were treated with MLi-2 (100 nM, 20 min) and then analyzed by immunoblot for phosphoRab10 and phosphoRab12 levels (Fig. 2; Supp. Figs. 1 and 2).

**Figure 2.**
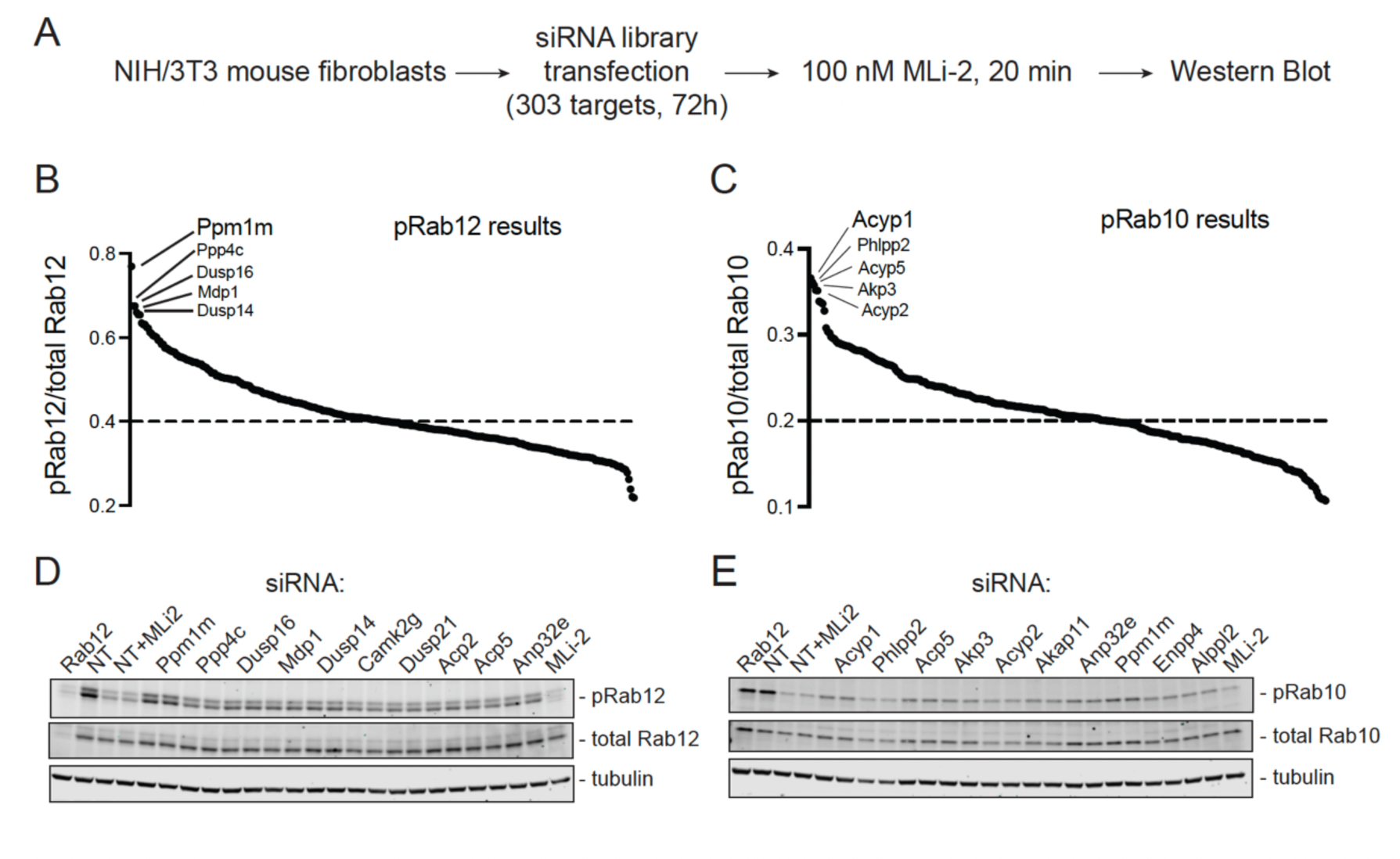
Phosphatome-wide siRNA screen in 3T3 cells reveals PPM1M as a phosphoRab12-preferring phosphatase. **(A)** Schematic describing screen workflow. **(B, C)** Summary plot of (B) pRab12/total Rab12 or (C) pRab10/total Rab10 levels after 72h siRNA and 20 min MLi-2 treatment, normalized to non-targeting control. Top 5 hits for each pRab as indicated. **(D, E)** Repeat immunoblots of the lysates of the top 10 hits from (B) for pRab12 and (C) for pRab10.

Figures 2B and 2C summarize the results of the screen, monitoring phosphoRab12 (Fig. 2B) and phosphoRab10 (Fig. 2C) levels, normalized to a non-targeting (NT) siRNA control. Genes that when knocked down, yielded the highest levels of a phosphoRab were considered the most significant LRRK2-counteracting phosphatases. The top 10 hits for each phosphoRab were re-analyzed to validate our findings (Fig. 2D,E). PPM1M was the top hit for phosphoRab12, and it was also among the top 10 hits for phosphoRab10. PPM1M is very closely related to PPM1H (∼45% identity) and is a member of the PPM1H subfamily of PPM phosphatases, along with PPM1J (*17*). Note that PPM1H was not identified in this screen to be a top hit for phosphoRab10, likely due to its absence or extremely low expression in the 3T3 cells used here; the prior phosphoRab10 phosphatase screen was carried out in A549 cells (*10*).

As expected, siRNA knockdown of Rab12 decreased total Rab12 but not total Rab10 levels. siRNA knockdown of Rab12 also decreased phosphoRab10 levels (normalized to total Rab10) because Rab12 activates LRRK2 Rab10 phosphorylation (*18, 19*). In additional controls, non-targeting siRNA samples treated with 100 nM MLi-2 for 2 hours also showed significant decreases in both phosphoRab10 and phosphoRab12 levels due to LRRK2 inhibition, confirming the activity of MLi-2. PPM1M knockdown in mouse embryonic fibroblasts (MEFs) also yielded an increase in phosphoRab12 levels, validating PPM1M’s role in a different cell line. In MEF cells, there was a slightly higher increase in phosphoRab12 levels upon PPM1M knockout than seen in 3T3 cells, likely due to higher LRRK2 expression (Supp. Fig. 3A,B).

### PPM1M overexpression influences phosphoRab12 and phosphoRab10 levels

Exogenous expression of PPM1M and PPM1H was used to assess the effects of these enzymes on phosphoRab10 and phosphoRab12 in cells. We also overexpressed PPM1H- and PPM1M-catalytically inactive mutants (PPM1H H153D and PPM1M H127D (*12*)) and substrate trapping mutants (PPM1H D288A and PPM1M D235A (*12*)) in HEK293 cells, along with LRRK2 R1441C (Fig. 3A). Note that HA-PPM1H and HA-PPM1M are expressed at different levels in these experiments, after transfection of cells with the same amount of transfected plasmid.

**Figure 3.**
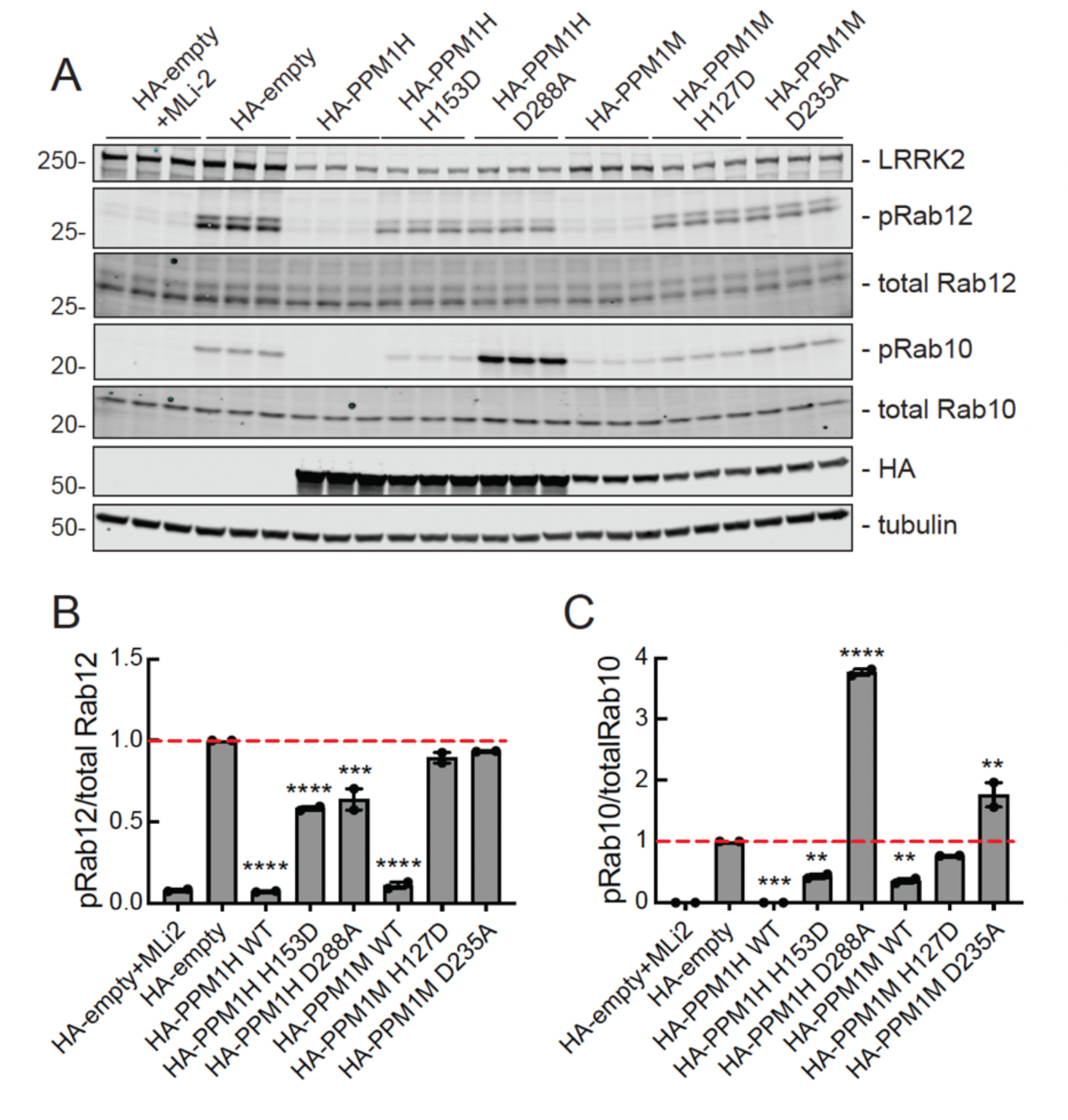
PPM1M overexpression preferentially decreases phosphoRab12 compared with phosphoRab10. **(A)** Immunoblot analysis of HEK293 cells overexpressing Flag-LRRK2 R1441C and HA-empty or HA-tagged PPM1H, PPM1H 153D, PPM1H D288A, PPM1M, PPM1M H127D, or PPM1M D235A. Cells were treated with 200 nM MLi-2 for 2h where indicated. **(B)** Quantitation of pRab12/total Rab12 and **(C)** pRab10/total Rab10 levels from immunoblots in (A), normalized to 1.0 for HA-empty. Error bars indicate SEM from two independent experiments analyzed in duplicate. Statistical significance determined by one way ANOVA, relative to HA-empty. For pRab10, ***p=0.0002 for PPM1H WT, **p=0.0097 for PPM1H H153D, ****p<0.0001 for PPM1H D288A, **p=0.0044 for PPM1M WT, **p=0.0014 for PPM1M D235A. For pRab12, ****p<0.0001 for PPM1H WT, PPM1H H153D, PPM1M WT, ***p=0.0002 for PPM1H D288A.

As expected, exogenously expressed PPM1H fully dephosphorylated phosphoRab10 and mostly dephosphorylated phosphoRab12 (Fig. 3A, lanes 7-9). PPM1M almost completely dephosphorylated phosphoRab12 and mostly dephosphorylated phosphoRab10 (Fig. 3A, lanes 16-18). Thus, upon overexpression, PPM1H and PPM1M can both dephosphorylate phosphoRab12, but PPM1H dephosphorylates phosphoRab10 more efficiently than PPM1M.

Also as expected, catalytically inactive mutants PPM1H H153D and PPM1M H127D failed to dephosphorylate either phosphoRab substrate (Fig. 3). The PPM1H D288A substrate-trapping mutant increased phosphoRab10 levels about four-fold, as previously shown (*12*), likely by shielding phosphoRab10 from other phosphatases. The PPM1M D235A substrate-trapping mutant increased phosphoRab10 levels about two-fold (Fig. 3C). Neither substrate-trapping mutant increased phosphoRab12 levels, suggesting that phosphoRab12 may be unique in the way that it engages these phosphatases (Fig. 3B). In summary, overexpression experiments show that PPM1M can work on both phosphoRab10 and phosphoRab12 substrates, with a preference for phosphoRab12.

### PPM1M knockout confirms preference for phosphoRab12 in culture and in mouse tissue

We generated pooled CRISPR knockout MEF cells to obtain lines without *Ppm1h*, *Ppm1m*, or the related *Ppm1j* (Fig. 4A). Due to the lack of a PPM1M or PPM1J antibody, CRISPR knockout of the gene was confirmed by DNA sequencing (Supp. Fig 3C). Of the three phosphatases, only the absence of PPM1M increased phosphoRab12 levels (Fig. 4A,B). For phosphoRab10, *Ppm1h* knockout increased its levels by almost two-fold as reported previously (*12*); *Ppm1m* knockout also increased phosphoRab10 levels by about 50% (Fig. 4B). *Ppm1j* knockout did not influence either phosphoRab10 or phosphoRab12 (Fig. 4B), and PPM1J has not shown activity on any phosphoRab tested to date (this study and (*12*)). It is possible that PPM1J is not expressed in these cells.

**Figure 4.**
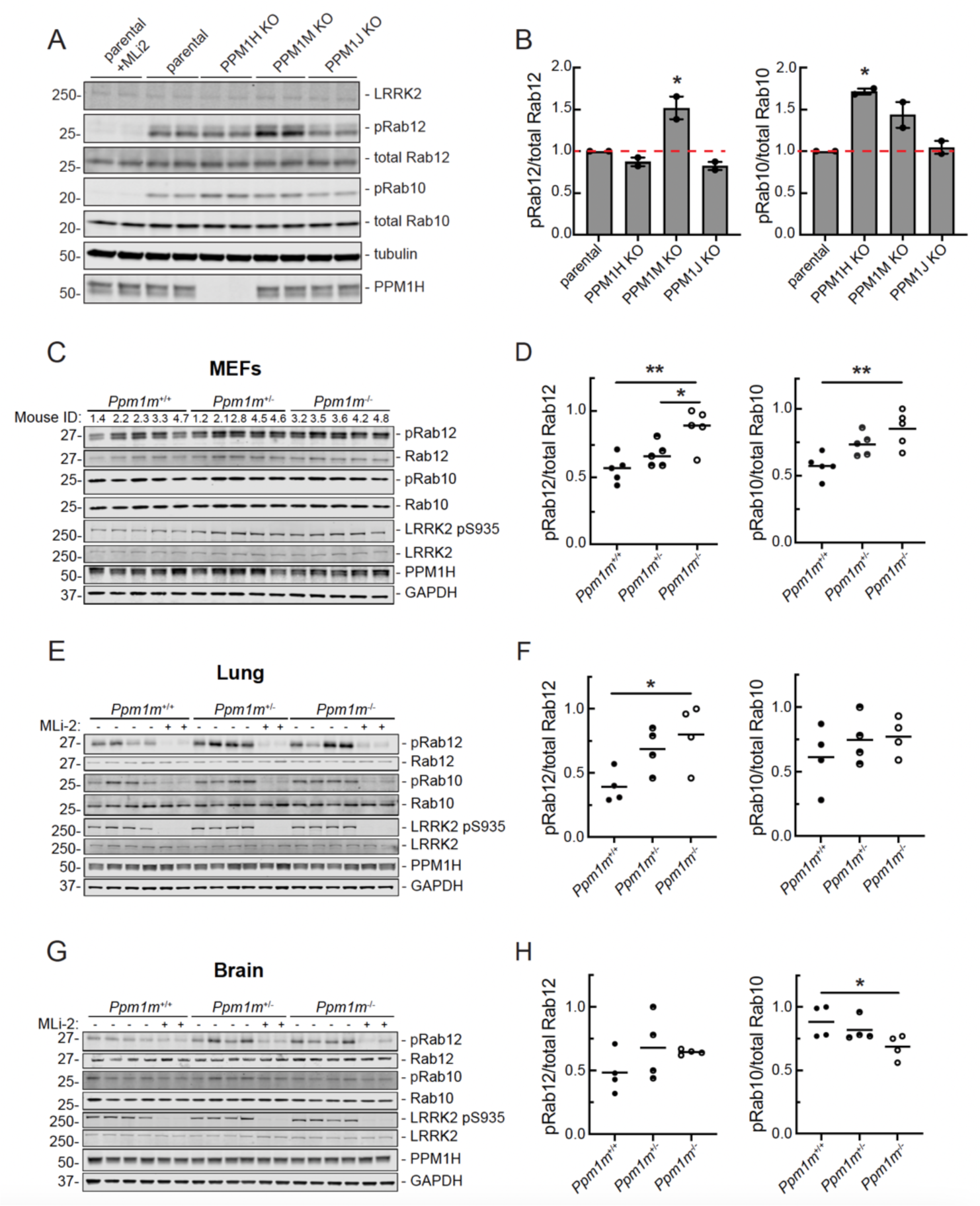
Knockout of PPM1 subfamily phosphatases in MEF cells and tissues confirms PPM1M substrate preferences. **(A)** Immunoblot analysis of parental (wild-type) and PPM1H, PPM1M, and PPM1J pooled knockouts in MEF cells. **(B)** Quantitation of pRab12/total Rab12 and pRab10/total Rab10 levels from immunoblots in (A), normalized to 1.0 for parental. Error bars indicate SEM from two independent experiments analyzed in duplicate. Statistical significance determined by one way ANOVA, respective to parental. For pRab10, *p=0.0122 for PPM1H knockout. For pRab12, *p=0.0117 for PPM1M knockout. **(C)** Immunoblot analysis of mouse embryonic fibroblasts (MEFs) derived from *Ppm1m* wild-type (*Ppm1m*^+/+^), heterozygous knockout (*Ppm1m*^+/-^), or homozygous knockout (*Ppm1m*^-/-^) mice. **(D)** Quantitation of pRab12/total Rab12 and pRab10/total Rab10 levels from immunoblots in (C), normalized to 1.0 for the highest value. Each dot represents the average of two independent replicates from one mouse. Statistical significance determined by one way ANOVA. For pRab10, **p=0.0036, for pRab12 *p=0.0399 and **p=0.0029. **(E)** Immunoblot analysis of lung lysates from mice as in (C). **(F)** Quantitation of immunoblots in (E), normalized as in D. Each dot represents the average of three independent replicates from one mouse. For pRab12, *p=0.0335. **(G)** Immunoblot analysis of whole brain lysates from mice as in (C). **(H)** Quantitation from immunoblots in **(G)**, as in F. Statistical significance determined by Kruskal-Wallis test for pRab10 and one way ANOVA for pRab12. For pRab10, *p=0.0349.

Similar experiments using MEF cells derived from homozygous *Ppm1m* knockout mice showed increases in phosphoRab12 and phosphoRab10 levels (Fig. 4C,D). Lung lysates from *Ppm1m* knockout mice showed an increase in phosphoRab12 levels with no change in steady state phosphoRab10 levels (Fig. 4E,F). A similar trend of increased phosphoRab12 levels was observed in *Ppm1m* knockout whole brain lysates but did not reach significance (Fig. 4G,H). The mouse knockout data are consistent with the general preference of PPM1M for phosphoRab12 over phosphoRab10 in cultured cells and in tissues. The *Ppm1m* knockout mouse exhibits no overt phenotype under standard husbandry conditions. Body weight, growth, and overall health appear normal, with no significant differences compared to wild-type littermates.

### Characterization of PPM1M

To further characterize PPM1M’s substrate preferences, we performed *in vitro* phosphatase assays using purified PPM1M and phosphoRab10 and phosphoRab12 (Supp Fig. 4A) in conjunction with phosphoRab-specific antibodies and immunoblotting to monitor phosphatase action (Fig. 5A). PPM1M displayed an approximately two-fold preference for phosphoRab12 over phosphoRab10; dephosphorylation rates were measured at different enzyme concentrations (Fig. 5B). We were not able to purify sufficient phosphorylated Rab12 and active PPM1M to determine precise k_cat_/K_m_ values by colorimetric assays. Also note that unlike PPM1H, the PPM1M enzyme lost significant activity upon freezing.

**Figure 5.**
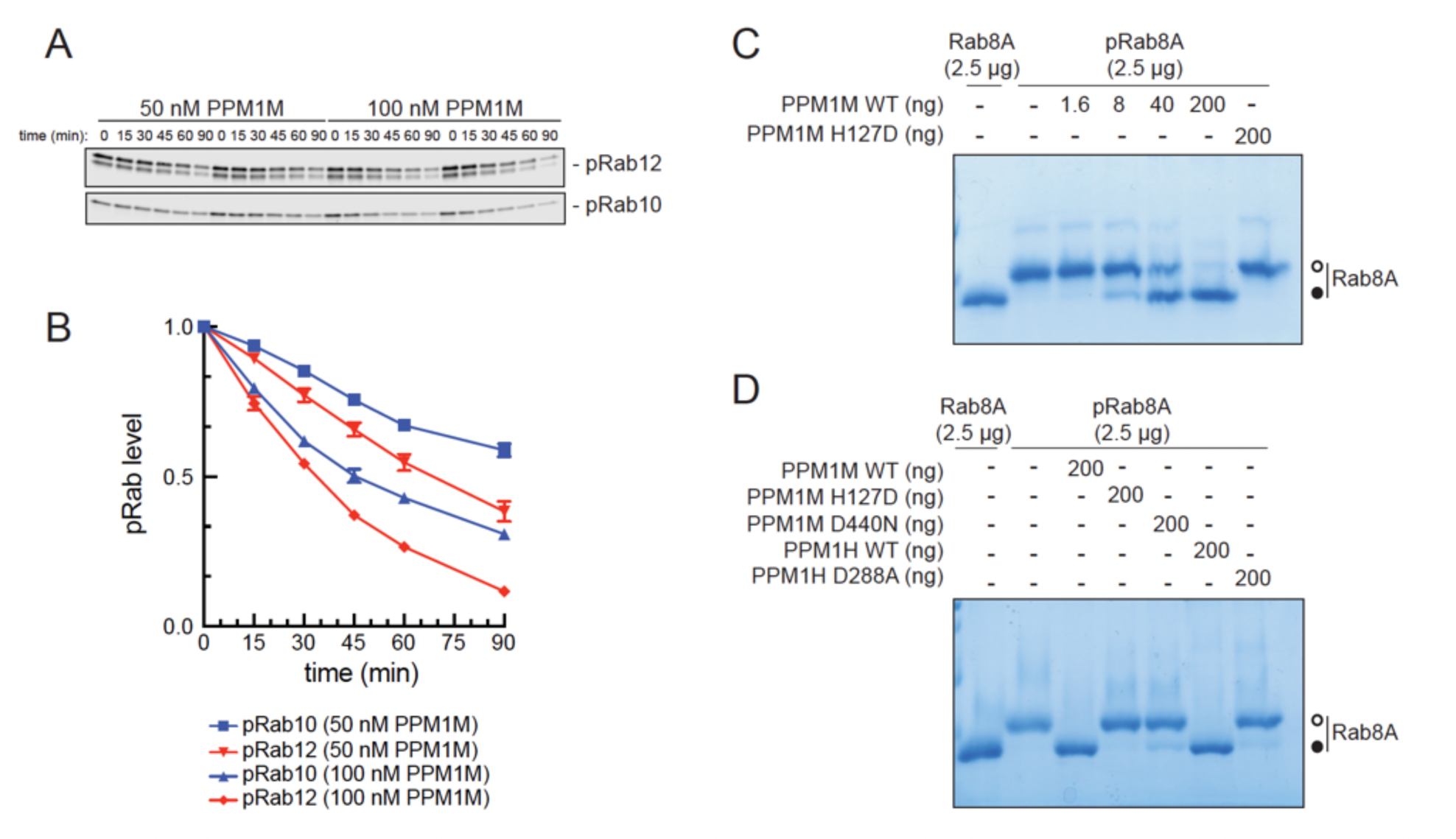
PPM1M prefers phosphoRab12 over phosphoRab10 *in vitro*. **(A)** Immunoblot analysis of pRab12 and pRab10 levels from *in vitro* biochemical reactions using 1.5μM pRab12 or pRab10 substrate and 50 or 100nM PPM1M enzyme, as indicated. **(B)** Quantitation of pRab12 and pRab10 levels from immunoblots in (A), normalized to 1.0 for 0 min. Error bars represent SEM from five independent experiments. **(C)** Phos-tag gel analysis of pRab8A dephosphorylation after *in vitro* biochemical reactions containing 2.5 µg pRab8A and indicated amounts of PPM1M. **(D)** Phos-tag gel analysis of pRab8A dephosphorylation after *in vitro* biochemical reactions containing 2.5 µg pRab8A and 200 ng of the indicated phosphatases.

Phos-tag gels used to resolve phosphorylated and unphosphorylated Rab8A showed that PPM1M can also act on phosphoRab8A at similar ratios of enzyme-to-substrate employed in panels 5A and 5B (Fig. 5C). In these experiments, PPM1M was as active as PPM1H in dephosphorylating Rab8A protein, while the catalytically inactive mutants (PPM1M H127D, PPM1H 153D) showed no activity (Fig. 5D).

PPM1H is a dimer (*16*). When PPM1M was exogenously expressed in HEK293 cells and the cytosol fractionated by size exclusion column chromatography, immunoblotting of column fractions indicated that PPM1M also exists as a dimer, based on its chromatographic properties in relation to marker proteins (Supp Fig. 4B).

### PPM1M flap domain contributes to its substrate selectivity

Elegant domain swap experiments have shown that the flap domain, located adjacent to the enzyme active site and unique to PPM family phosphatases, is critical for substrate specificity (Fig. 6A, navy blue; (*16*)). Similar domain swaps between the flaps of PPM1H and PPM1M confirmed their roles in phosphoRab selectivity (Fig. 6B-D). Constructs were expressed in HEK293 cells in the presence of LRRK2 R1441C to compare phosphoRab12 and phospho-Rab10 levels (Fig. 6C,D). As expected, PPM1H preferentially dephosphorylated phosphoRab10 over phosphoRab12, while PPM1M dephosphorylated phosphoRab12 more than phospho-Rab10. PPM1H containing the PPM1M flap domain (PPM1H_M flap) no longer dephosphorylated phosphoRab10 as efficiently, yet it retained a similar—if not improved—ability to work on phosphoRab12. PPM1M containing PPM1H’s flap domain PPM1M_H flap) was expressed at about two-fold lower levels than wild-type PPM1M. Furthermore, it was unable to dephosphorylate phosphoRab12 to the same extent as the parental PPM1M protein, despite dephosphorylating phosphoRab10 to a similar extent as PPM1M (Fig. 6D). Therefore, PPM1H and PPM1M both require their flap domains for proper substrate recognition.

**Figure 6.**
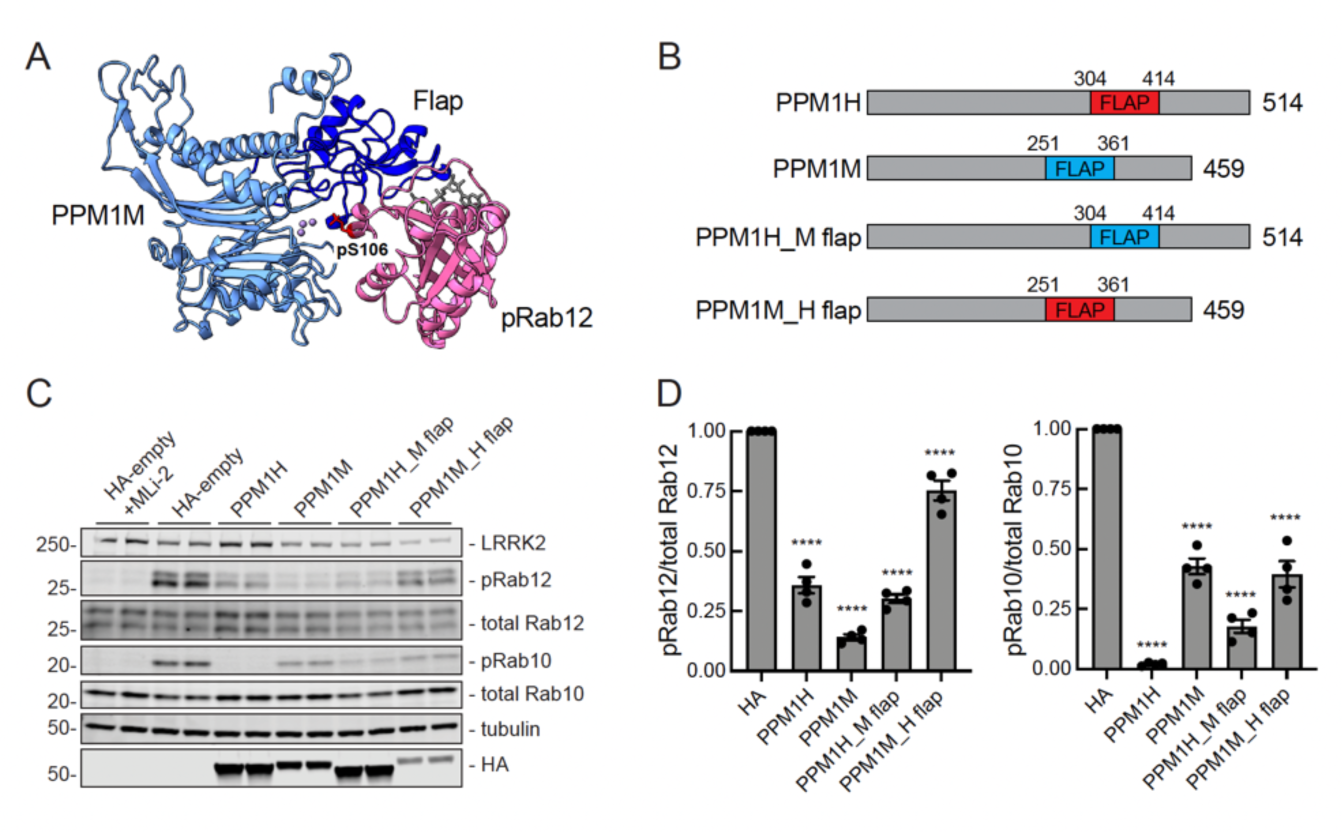
PPM1H and PPM1M flap domains are necessary for proper substrate recognition. **(A)** Alphafold modeling of PPM1M (blue) and pRab12 (magenta). The PPM1H flap domain is shown in navy; phosphoserine 106 of the pRab12 substrate is indicated at the metal-containing PPM1M active site. **(B)** Diagram of PPM1H and PPM1M swapped flap domain constructs. **(C)** Immunoblot analysis of HEK293 cells overexpressing Flag-LRRK2 R1441C and HA-empty or HA-tagged PPM1H, PPM1M, PPM1H with PPM1M flap domain (PPM1H_M flap), or PPM1M with PPM1H flap domain (PPM1M_H flap). **(D)** Quantitation of pRab12/total Rab12 and pRab10/total Rab10 levels from immunoblots in (C), normalized to 1.0 for HA-empty. Error bars indicate SEM from four independent experiments analyzed in duplicate. Statistical significance determined by one way ANOVA, respective to HA-empty, ****p<0.0001.

### *Ppm1m* knockout phenocopies hyperactive *Lrrk2* knock-in ciliation defect

We have shown previously that mice harboring hyperactive *Lrrk2* mutations (R1441C or G2019S) or lacking *Ppm1h* show loss of primary cilia in cholinergic interneurons and astrocytes of the mouse dorsal striatum (*20–22*). Similar cilia losses are seen in dorsal striatum from patients with idiopathic or LRRK2-pathway mutations (*22*). Therefore, we examined the ciliation status of cholinergic interneurons of the dorsolateral striatum of mice lacking *Ppm1m*. As shown in Fig. 7, while cholinergic interneurons were ∼70% ciliated in the dorsal striatum of wild-type mice, they were only ∼45% ciliated in *Ppm1m* knockout mice. As expected (*20, 21*), there was no change in ciliation in the surrounding neurons, most of which are medium spiny neurons. Thus, at least in cells of the dorsal striatum, PPM1M plays as significant a role in regulating the LRRK2 pathway as PPM1H or pathogenic LRRK2.

**Figure 7.**
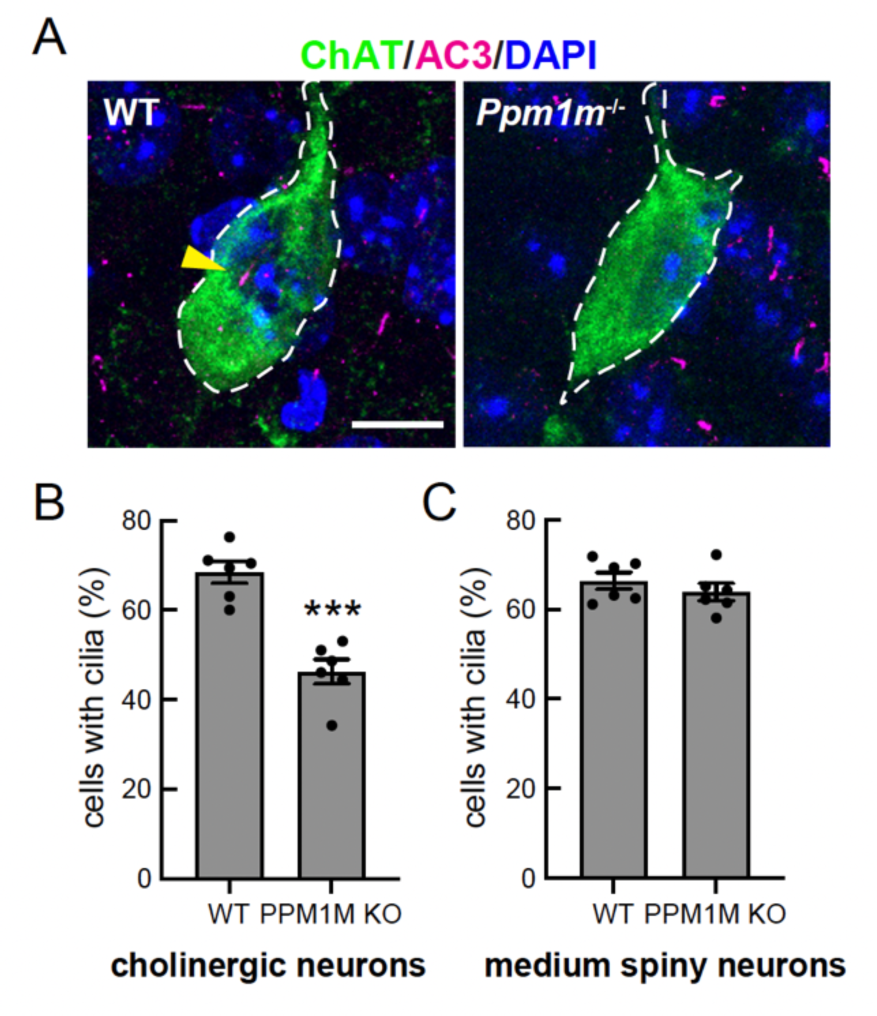
*PPM1M* knockout phenocopies *LRRK2* hyperactive ciliation phenotype. **(A)** Example confocal immunofluorescence micrographs of sections of the dorsal striatum from 3-month-old wild-type or *Ppm1m*^-/-^ mice; scale bar, 10 μm. Cholinergic interneurons were labeled using anti-choline acetyltransferase (ChAT) antibody (green); primary cilia were labeled using anti-AC3 (adenylate cyclase 3) antibody (magenta; yellow arrowhead). Nuclei were labeled using DAPI (blue). **(B)** Quantitation of ChAT^+^ neurons and **(C)** surrounding (mostly medium spiny) neurons containing a cilium. Error bars represent SEM from six individual brains per group, two to three sections per mouse. >36 ChAT^+^ neurons and >500 ChAT^-^ cells were scored per mouse. Statistical significance was determined using an unpaired t-test. ***p=0.0001 for cholinergic neurons; ns p=0.3835 for medium spiny neurons.

### *PPM1M* p.D440N mutant increases risk of Parkinson’s disease

Mutations in *LRRK2* and *PPM1H* are linked to PD. Therefore, we explored the possibility of a similar genetic link between *PPM1M* mutations and PD. Our recent studies identified *RAB32* p.R71R as a susceptibility variant for familial PD (*23–25*) and also reported additional variants in other genes with strong associations with PD (*25*). Interestingly, among them was a rare missense variant in the *PPM1M* gene: NM_144641.4:c.1318G>A p.D440N (hereafter referred to as D440N). This variant was found in three out of 2,184 familial index patients with PD, with only three occurrences in a much larger control group of 69,775 subjects (odds ratio 44, p=3.81E-05). For one of the three index patients, DNA was available for additional family members, allowing us to test for segregation of the *PPM1M* D440N variant (Fig. 8A). The carrier status of the affected brother was inferred to be heterozygous for the *PPM1M* D440N variant, given that both children carried the variant. However, a second-degree cousin, who also had PD, did not carry the variant (Fig. 8A).

**Figure 8.**
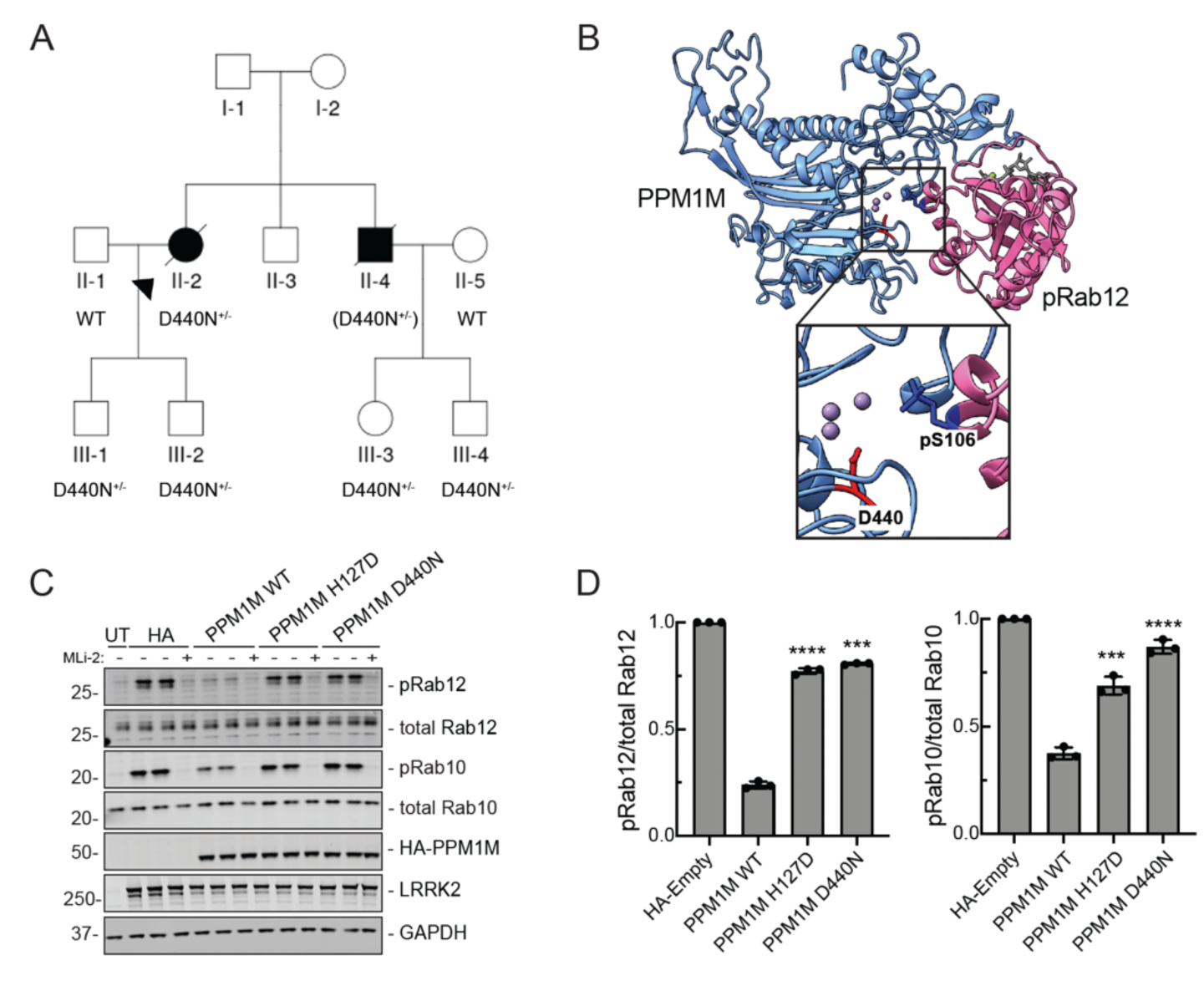
Parkinson’s disease patient linked to PPM1M D440N mutation. **(A)** Individuals genotyped for the c.1318G>A p.D440N mutation are indicated as D440N^+/-^. In the case of II-4, the mutation was inferred from its presence in both of his children, III-3 and III-4, and is shown in brackets (D440N^+/-^). A second-degree cousin of II-3 and II-4 also had PD but did not carry the D440N variant. The shared common ancestors of this individual with II-2 and II-4 are their great-grandparents (the grandparents of I-2, not shown in the pedigree). No clinical phenotype information for any of the members of generation III was available. **(B)** Alphafold modeling of D440 in the PPM1M (blue) active site shown with pRab12 (magenta, residues 38-222). Inset shows enlarged view of PPM1M D440 and Rab12 pS106. **(C)** Immunoblot analysis of HEK293 cells overexpressing Flag-LRRK2 R1441G and HA-empty, HA-PPM1M WT, HA-PPM1M H127D, HA-PPM1M D440N, with untransfected (UT) control. MLi-2 (200 nM) treatment was for 90 min as indicated. **(D)** Quantitation of pRab12/total Rab12 and pRab10/total Rab10 levels from immunoblots in (C), normalized to HA-empty. Error bars indicate SD from three independent experiments carried out in duplicate. Statistical significance determined by Welch’s t-test, followed by Benjamini-Hochberg correction for multiple comparisons, respective to HA-empty. ***p=0.00011 for pRab12 - PPM1M D440N, ***p=0.00078 for pRab10 - PPM1M H127D, otherwise ****p<0.0001.

We also identified the *PPM1M* D440N variant in one out of 382 patients from our Austrian cohort of PD patients. This patient had early onset PD (age 43) but no known family history of PD. The *PPM1M* D440N variant was also identified in three additional subjects from other PD cohorts: once in 700 cases from the Mayo Clinic’s brain bank in a subject with dementia with Lewy bodies; once in 725 PD cases from a Polish cohort; and once in 139 Ukrainian PD cases. However, we did not identify this variant in 10,000 subjects with PD from the Centogene database where exome or genome data was available (Supp. Table 3). Overall, we detected the *PPM1M* D440N variant in seven of 14,835 individuals with PD or PD-related conditions (minor allele frequency: 2.36 x10E-04). In control databases, the frequency is approximately six times lower: gnomAD (4.13E-05) and Centogene (3.57E-05).

Given the rarity of the *PPM1M* D440N variant, we were unable to statistically confirm *PPM1M* as a novel PD gene. However, we have obtained additional functional evidence that *PPM1M* D440N may contribute to PD via indirect activation of the LRRK2 kinase pathway.

### PPM1M D440N is inactive

The *PPM1M* D440N mutation is located precisely in the middle of PPM1M’s active site (Fig. 8B). Therefore, we tested if this mutation affects PPM1M phosphatase activity. Upon expression in HEK293 cells, the PPM1M D440N mutant protein was inactive, comparable to the PPM1M phosphatase-dead H127D mutation (Fig. 8C,D). Similarly, in vitro dephosphorylation assays with purified proteins, visualized using Phos-tag gels, showed that PPM1M D440N is as inactive as PPM1M H127D on phosphoRab8A (Fig. 5D). Future work will be needed to further characterize the possible contribution of the *PPM1M* D440N mutation to PD.

## Discussion

We have shown here that like PPM1H, the related PPM1M protein is a Rab-phosphatase that counteracts LRRK2 phosphorylation. Although both phosphatases can act on phosphoRab8A and phosphoRab10, PPM1M is unique in its ability to act on phosphoRab12 at endogenous levels in 3T3 and MEF cells. In MEFs from a mouse *Ppm1m* knockout model, we detect increased phosphoRab12 and phosphoRab10 levels. Similar results were obtained when comparing the relative consequences of *Ppm1m* and *Ppm1h* CRISPR knockout in wild-type MEF cells. In lung lysates, we found that *Ppm1m* knockout increases phosphoRab12 but not phosphoRab10 levels, with a similar trend in whole brain lysates.

Using purified phosphatases and phosphorylated Rab proteins, PPM1M shows a roughly two-fold preference for phosphoRab12 over phosphoRab10. Both PPM1H and PPM1M are expressed at low levels in most cells tested (approximately 10,000 copies per MEF cell), and the lack of available anti-PPM1M antibodies has made it challenging to study endogenous PPM1M. Nevertheless, both PPM1M and PPM1H’s roles as Rab phosphatases are now established through knockout experiments in both cell culture and mouse models. Consistent with previous reports (*12*) we were unable to detect any activity for the related PPM1J protein.

Alphafold modeling based on the crystal structure of PPM1H (*16*) reveals that PPM1M shares a conserved phosphatase fold. The model also indicates that PPM1M has a similar overall structure, but with a smaller, substrate-specifying flap domain. In general agreement with previously reported domain swap experiments, the flap domain of PPM1H inserted in place of PPM1M’s flap domain decreases PPM1M’s ability to act on phosphoRab12. Similarly, insertion of the PPM1M flap into PPM1H slightly decreases its ability to dephosphorylate phosphoRab10. Although neither swap was perfect, these data support the conclusion that the flap domain is an important contributor to Rab substrate specificity of PPM family phosphatases.

PPM1H relies on an N-terminal amphipathic helix to bind highly curved liposomes in vitro and to localize approximately half of the protein to the Golgi in cultured cells (*14*). In contrast, PPM1M lacks this N-terminal amphipathic helix, consistent with our observation that exogenously expressed PPM1M appears entirely cytosolic in 3T3 and A549 cells. Like PPM1H (*16*), PPM1M forms a dimer in the cytosol, as determined by gel filtration, and the protein maintains its dimeric state after purification from bacteria. Due to low PPM1M levels, we cannot rule out the possibility that a fraction of the endogenous protein is membrane-associated. We have obtained evidence for heterodimers when analyzing overexpressed PPM proteins, but these likely reflect a small proportion of the total enzyme pool.

PPM1H is widely expressed across various tissues, with the highest levels in the brain, whereas PPM1M is generally expressed at lower levels and is most abundant in neutrophils. Thus, the two proteins diverge in their expression patterns. Notably, while PPM1H is highly expressed in the brain, its levels vary significantly among different neuronal cell types. For example, in the striatum, we have shown that medium spiny neurons exhibit high LRRK2 and low PPM1H expression (*22*). Cholinergic interneurons, though, show the opposite pattern, with low LRRK2 and high PPM1H expression (*22*). Despite the apparently low expression in the brain, we detected increased levels of phosphoRab12 in brain extracts from PPM1M knockout mice. PPM1M knockout mice also show loss of primary cilia in cholinergic neurons of the dorsal striatum, much like mice harboring hyperactive LRRK2 or PPM1H inactivating mutations. The consequences of increased phosphoRab12 in the brains of these mice will be an important area for future investigation.

We identified a *PPM1M* variant that was reported to increase risk of PD, D440N (*25*). Large-scale screening of available genetic data identified a number of PD patients that carry the *PPM1M* D440N variant, and the occurrence of this variant is at least ∼6-fold higher in PD cohorts compared with extremely large control data sets. Determining the pathogenicity of rare variants is a growing challenge within the field of clinical genetics. The small number of carriers precludes statistical evidence to confer a designation of effect, and thus functional assays will be critical in determining pathogenicity. The PPM1M D440N mutation is located precisely in the phosphatase active site. Upon expression in cultured cells or after purification from bacteria, PPM1M D440N protein showed complete loss of activity, supporting functional pathogenicity. Integration of functional data with further genetic screening efforts, both at the variant and gene level, will help resolve the role of *PPM1M* variants in PD susceptibility.

Despite the strong connection between PPM1H and PPM1M phosphatases and LRRK2 action, additional phosphatases are also clearly involved in regulating overall phosphoRab levels. Moreover, phosphoRab12 seems more resistant to phosphatases than phosphoRab10 based on dephosphorylation kinetics in A549, HEK293, 3T3, and MEF cells. Part of this resistance may be because Rab12 is also phosphorylated by additional kinases. Nevertheless, the screens presented herein and the prior Berndsen et al. (*12*) screen have identified a group of additional phosphatases with the capacity to act on phosphorylated Rab proteins. Additional phosphatases are especially relevant given the findings of Ito et al. (*26*) who have shown that Rab phosphorylation is a highly transient modification. Their data suggest that overall phosphorylation is regulated by a constant balance of kinase activation countered by dephosphorylation. Future experiments will provide additional insight into the link between LRRK2-dependent Rab phosphorylation and phosphatase action.

## Materials and Methods

### Cloning and plasmids

DNA constructs were amplified in *E. coli* DH5α and purified using mini prep columns (Econospin, Fisher Scientific). Whole plasmid DNA sequence verification was performed by Primordium/Plasmidsaurus (https://plasmidsaurus.com). pET15b His-MST3 was a kind gift of Amir Khan (Trinity College, Dublin). pCMV5D HA-empty or HA-PPM1H, PPM1H H153D, PPM1H D288A, PPM1M, PPM1M H127D, PPM1M D235A, and PPM1M D440N were obtained from MRC PPU. pET15b His-SUMO-Rab10 Q68L and Rab12 Q101L were previously obtained or cloned. pET15b His-SUMO-PPM1M was cloned using the pET15b backbone and HA-PPM1M insert via Gibson assembly. A detailed protocol for Gibson assembly can be found here https://doi.org/10.17504/protocols.io.eq2lyjwyqlx9/v1. See Supplemental Table 1 (Key Resource Table) for access to plasmids from MRC PPU and/or Addgene.

### Cell culture

HEK293 cells were purchased from ATCC. Flp-In 3T3 cells were purchased from Invitrogen (Carlsbad, CA). PPM1H knockout A549, wild-type MEF, and PPM1M knockout MEF cells were obtained from MRC-PPU. All cells were cultured in DMEM high-glucose media (Cytiva, Marlborough, MA) with 10% fetal bovine serum (Sigma, St. Louis, MO) and 1% Penicillin-Streptomycin (Sigma) and grown at 37°C, 5% CO_2_ in a humidified atmosphere and regularly tested for *Mycoplasma* contamination.

### siRNA phosphatase screen

Dharmacon mouse phosphatase siRNA library was purchased from Horizon Discovery (Cambridge, UK). This library does not contain all phosphatase genes; therefore a cherry-picked custom library of additional genes was also purchased (see Supplemental Table 2) from Horizon Discovery. Low passage 3T3 Flp-In mouse fibroblasts were plated in 6-well plates at ∼30% confluency (200,000 cells/well) in 1.6 mL complete DMEM, 16-24 hours prior to transfection. siRNA resuspension in Dharmafect 1 transfection reagent (Dharmacon, Lafayette, CO) was performed according to the manufacturer. In brief, siRNA was resuspended in 1x siRNA buffer (Dharmacon) to a final concentration of 20 µM. Each well was transfected with 25 nM siRNA and 4 µL Dharmafect 1 in a total of 400 µL Opti-MEM. Transfection mixture was added dropwise to each well. After 24 hours, transfection media was replaced with fresh media, and cells were maintained for another 48 hours. 100 nM of MLi-2 was then added to each well for 20 minutes prior to harvesting. In some control samples, 100 nM MLi-2 was added to cells for 2 hours. Cells were harvested, lysed, and analyzed by immunoblotting. See detailed protocol here: dx.doi.org/10.17504/protocols.io.36wgqdxr5vk5/v1.

### Transient overexpression in HEK293 cells

For transient overexpression assays in HEK293 cells, cells were plated in 2 mL complete media in 6-well plates 16-24 hours prior to transfection to achieve ∼75% confluency at the time of transfection. Plasmids were mixed in 200 µL Opti-MEM (Gibco, Grand Island, NY) with 1 mg/mL PEI (polyethylenimine; 1:5 DNA:PEI ratio) and allowed to incubate for 15 minutes at room temperature. 0.5 µg of all plasmids were used except for 1 µg Flag-LRRK2 R1441C. For experiments involving overexpression of PPM1M D440N (Figure 8), plasmids were mixed in 300 µL Opti-MEM (Gibco) with 1 mg/mL PEI (1:3 DNA:PEI ratio) and incubated for 30 minutes at room temperature. 0.75 µg of all plasmids were used except for 1.25 µg Flag-LRRK2 R1441G. Transfection mixture was added dropwise to attached cells and incubated for 24 hours prior to immunoblotting analysis.

### Pooled CRISPR knockout

Pooled CRISPR knockouts were generated by electroporation using guide RNAs designed and synthesized from Synthego. A detailed protocol can be found here: dx.doi.org/10.17504/protocols.io.bp2l6dqodvqe/v1. In brief, the top two ranked sgRNA per gene as determined by Synthego Knockout Design Tool (https://design.synthego.com) were pooled and combined with Cas9 (IDT) at a 6:1 sgRNA:Cas9 ratio to form a RNP complex. This RNP complex was electroporated into cells resuspended in Opti-MEM using a 0.2 cm cuvette and NEPA21 Type II electroporator. Cells were allowed to recover and expand for one week. Cells were then genotyped by extracting genomic DNA from cell pellets using QuikExtract and PCR amplification and sequencing of the targeted regions. Sequencing results were analyzed using Synthego ICE analysis (https://ice.synthego.com). Knockout was also validated by immunoblotting if endogenous protein levels were detectable (for PPM1H only, not PPM1M or PPM1J).

### Mice

Mice were generated at the Baylor College of Medicine as part of the Baylor College of Medicine, Sanger Institute, and MRC Harwell (BaSH) Consortium for the NIH Common Fund program for Knockout Mouse Production and Cryopreservation (1U42RR033192-01) and Knockout Mouse Phenotyping (1U54HG006348-01). Mice are distributed by MRC Harwell on behalf of MMRRC. More information can be found here: https://www.informatics.jax.org/allele/MGI:5638564.

### Isolation of PPM1M knockout MEFs

Wild-type, heterozygous, and homozygous PPM1M knockout MEFs were isolated from littermate matched mouse embryos at day E12.5 resulting from crosses between heterozygous PPM1M knockout and wild-type mice using the protocol described in dx.doi.org/10.17504/protocols.io.eq2ly713qlx9/v1. Genotypes were verified via allelic sequencing and immunoblotting analysis. Cells were cultured in DMEM containing 10% (v/v) FBS, 2 mM L-glutamine, penicillin-streptomycin 100 U/mL, 1 mM sodium pyruvate, and 1x non-essential amino acid solution. Genotyping was performed by TransnetYX (https://www.transnetyx.com/) using the following probes: Ppm1m-1 WT and Ppm1m Tm2b as specified by the vendor.

### Mouse tissue homogenization, cell lysis and immunoblotting analysis

Quantitative immunoblotting analysis from cultured cell lysates was performed as described in dx.doi.org/10.17504/protocols.io.bsgrnbv6. Briefly, cell pellets were collected and lysed in lysis buffer (50 mM Tris–HCl pH 7.4, 150 nM NaCl, 1 mM EGTA, 2% glycerol, cOmplete EDTA-free protease inhibitor cocktail (Roche), PhosSTOP phosphatase inhibitor cocktail (Roche), 1 μg/ml microcystin-LR (Sigma), and 1% (v/v) Triton X-100). Lysates were clarified by centrifugation at 10,000 X g at 4°C for 10 min.

For mouse tissues, after dissection, tissues were homogenised using the Precellys Tissue Homogeniser in Tissue Grinding CKMix50-R tubes (P000922-LYSK0-A) with lysis buffer (50 mM Tris–HCl pH 7.4, 150 nM NaCl, 1 mM EGTA, 2% glycerol, cOmplete EDTA-free protease inhibitor cocktail (Roche), PhosSTOP phosphatase inhibitor cocktail (Roche), 1 μg/ml microcystin-LR (Sigma), and 1% (v/v) Triton X-100) at 6800 rpm for 3×20 seconds, with 30 seconds between each round of lysis. Following homogenisation, lysates were clarified by centrifugation at 10,000 X g at 4°C for 10 minutes.

Lysate protein concentration was determined by Bradford assay. Samples containing equal protein amounts were mixed with 5x SDS sample buffer (250 mM Tris-HCl, pH 6.8, 30% glycerol (v/v), 10% SDS (w/v), 0.1% bromophenol blue (w/v), 10% 2-mercaptoethanol (BME) (v/v)) and resolved on 4-20% precast gels (Bio-Rad) then transferred onto nitrocellulose membranes using the Transblot Turbo System (Bio-Rad). Membranes were blocked in 5% milk with TBST for 1 hour and incubated with specific primary antibodies overnight at 4°C.

See Supplemental Table 1 (Key Resource Table) for primary and secondary antibodies used and their dilutions. Primary antibodies were diluted in 3% BSA in Tris-buffered saline (200 mM Tris, 1.5M NaCl) with 0.1% Tween-20 (TBST) and detected using LI-COR IRdye labeled secondary antibodies (donkey anti-mouse 680/800, donkey anti-rabbit 680/800), diluted 1:10,000 in 5% milk in TBST. Membranes were scanned on the LICOR Odyssey DLx scanner. Images were saved as .tif files and analyzed using the gel scanning plugin in ImageJ.

### Protein purification

His-SUMO-PPM1M, His-SUMO-Rab10 Q68L, and His-SUMO-Rab12 Q101L were purified after expression in E. coli BL21 (DE3 pLys). A detailed protocol can be found at dx.doi.org/10.17504/protocols.io.bu7wnzpe. In brief, bacterial cells were grown at 37°C in Luria Broth and induced at A600 nm = 0.6 by the addition of 0.3 mM isopropyl-1-thio-β-d-galactopyranoside (IPTG) (Gold Biotechnology, St. Louis, MO) and harvested after growth for 18 hours at 18°C. The cell pellets were resuspended in ice-cold lysis buffer (50 mM HEPES, pH 8.0, 10% [vol/vol] glycerol, 500 mM NaCl, 10 mM imidazole, 5 mM MgCl_2_, 0.2 mM TCEP, 20 μM GTP, and cOmplete EDTA-free protease inhibitor cocktail (Roche). The resuspended bacteria were lysed by one passage through an Emulsiflex-C5 apparatus (Avestin) at 10,000 lbs/in^2^ and centrifuged at 40,000 rpm for 45 min at 4°C in a Beckman Ti45 rotor. Cleared lysate was filtered through a 0.2 µm filter (Nalgene) and passed over a HiTrap TALON crude 1 mL column (Cytiva). The column was washed with lysis buffer until absorbance values reached pre-lysate values. Protein was eluted with a gradient from 20 to 500 mM imidazole-containing lysis buffer. Peak fractions were analyzed by 4-20% SDS-PAGE to locate protein. If cleaving off His-SUMO tag (for Rab10 and Rab12), the eluate was cleaved overnight using homemade SUMO protease while dialyzing into a buffer containing 50 mM HEPES pH 8, 5% (vol/vol) glycerol, 150 mM NaCl, 5 mM MgCl_2_, 0.2 mM TCEP, and 20 μM GTP. The cleaved product was then further purified by gel filtration on Superdex 200 16/60 120 mL or Superdex 200 10/300 24mL size exclusions columns (Cytiva) using a buffer containing 50 mM HEPES pH 8, 5% (vol/vol) glycerol, 150 mM NaCl, 5 mM MgCl_2_, 0.2 mM TCEP, and 20 μM GTP. Fractions were resolved on a 4-20% Mini-PROTEAN TGX Gel (Bio-Rad) and visualized using InstantBlue Coomassie Stain (Abcam, Waltham, MA).

### In vitro phosphatase assay with immunoblotting analysis

A detailed protocol can be found at dx.doi.org/10.17504/protocols.io.5jyl8d4j7g2w/v1. Briefly, untagged Rab10 Q68L or untagged Rab12 Q101L was incubated with His-MST3 kinase in a reaction buffer (50 mM HEPES pH 8,100 mM NaCl, 5 mM MgCl_2_,100 μM GTP, 0.5 mM TCEP, 10% glycerol, 5 μM BSA) at 4°C overnight to phosphorylate Rab10/Rab12. Next, His-MST3 kinase was removed by passing the sample through a 1-mL syringe column containing 100 μL (50%) Ni-NTA slurry; the flow through containing phosphorylated Rab10 or Rab12 was collected. For phosphatase assays, 1.5 µM pRab10 or pRab12 was incubated with 50 or 100nM His-SUMO-PPM1M at 30°C for various times. A master mix reaction was made using 15 µL total reaction volume per time point. At the indicated time point, 15 µL was removed from the reaction tube and reaction aliquots were stopped by addition of 5 µL SDS-PAGE sample buffer. Samples were then analyzed by immunoblotting to detect dephosphorylation of Rab10 or Rab12 using anti-pRab10 antibody or anti-pRab12 antibody as above.

### Chromatography of HEK293 cytosol

Crude membrane fractionation was performed according to https://doi.org/10.17504/protocols.io.yxmvmnb99g3p/v1. Briefly, 70% confluent HEK293 cells were transfected in 10 cm dishes with 5 µg HA-PPM1M plasmid. 24 hours after transfection, 3×10 cm dishes of cells were washed 2x with ice-cold PBS, pooled, and swollen in 400 µL of hypotonic buffer (10 mM HEPES pH 7.4 with protease inhibitors). After 20 min, 100 µL of 5X resuspension buffer was added to achieve a final concentration of 1X resuspension buffer (50 mM HEPES pH 7.4, 150 mM NaCl, 5 mM MgCl_2_, 1X cOmplete protease inhibitor cocktail (Roche), 1X phosphatase PhosStop inhibitor (Roche)). The suspension was passed 40 times through a 27G needle. Lysate was spun at 1,000 X g for 5 min at 4°C to pellet nuclei. The postnuclear supernatant was spun at 55,000 RPM for 20 min at 4°C in a tabletop ultracentrifuge in a TLA100.2 rotor; the resulting supernatant (∼500 µL) was collected as the cytosol fraction. The supernatant was then applied onto a 24 mL Superdex 10/300 column (Cytiva) and fractions subjected to immunoblot analysis to determine PPM1M elution. Corresponding mass (kDa) was determined by comparison with calibration standards (Bio-Rad).

### In vitro pRab8A phosphatase assays with Phos-tag gel analysis

Bacterial expression and purification of 6xHIS-SUMO-PPM1M (WT, H127D, D440N), PPM1H (WT, D288A) and large scale preparation of phosphorylated GTPγS-bound Rab8A Thr72 were performed as previously described (https://doi.org/10.17504/protocols.io.bu7wnzpe and dx.doi.org/10.17504/protocols.io.butinwke). For in vitro phosphatase assays, 2.5 µg phosphorylated, GTPγS-bound Rab8A was incubated with varying amounts of recombinant wild-type PPM1M or PPM1H or their respective variants in a reaction buffer containing 40 mM HEPES (pH 7.0) and 10 mM MgCl_2_. GTPγS-bound pRab8A was in 20 mM MES pH 5.3, 0.1 M NaCl, 10% (by vol) glycerol, 0.03% (by vol) Brij 35, 14 mM 2-mercaptoethanol, 2 mM MgCl_2_, 1 mM GTPγS while PPM1M/H proteins were in a buffer containing 50 mM Tris/HCl pH 7.5, 150 mM NaCl, 2 mM MnCl_2_, 0.5 mM TCEP buffer. Following a 30-minute incubation, reactions were terminated by adding 6uL of 4x lithium dodecyl sulfate (LDS) (106 mM Tris HCl, 141 mM Tris Base, 2% (w/v) LDS, 10% (v/v) glycerol, 0.51 mM EDTA, 0.22 mM SERVA Blue G250, 0.175 mM Phenol Red, pH 8.5) supplemented with 5% (v/v) 2-mercaptoethanol. Samples were analysed via Phos-Tag gel electrophoresis as described by Ito et al. (*16*). Post-electrophoresis, the gel was stained with Instant Blue Coomassie (Abcam) for subsequent analysis.

### Mouse brain processing

Ppm1m^-/-^ (3-month-old) mice and their littermate wild-type controls were fixed by transcardial perfusion using 4% paraformaldehyde (PFA) in PBS as described in dx.doi.org/10.17504/protocols.io.bnwimfce. Whole brain tissue was extracted, post-fixed in 4% PFA for 24 hr and then immersed in 30% (w/v) sucrose in PBS until the tissue settled to the bottom of the tube (∼48 hours). The brains were harvested in Dundee and sent to Stanford with identities blinded until analysis was completed. Prior to cryosectioning, brains were embedded in cubed-shaped plastic blocks with OCT (BioTek, USA) and stored at −80°C. OCT blocks were allowed to reach −20°C for ease of sectioning. The brains were oriented to cut coronal sections on a cryotome (Leica CM3050S, Germany) at 25 μm thickness and positioned onto SuperFrost plus tissue slides (Thermo Fisher, USA).

### Immunohistochemical staining

Mouse brain striatum was subjected to immunostaining following a previously established protocol (dx.doi.org/10.17504/protocols.io.bnwimfce). Frozen slides were thawed at room temperature for 15 minutes and then gently washed twice with PBS for 5 minutes each. Antigen retrieval was achieved by incubating the slides in 10 mM sodium citrate buffer pH 6.0, preheated to 95°C, for 15 minutes. Sections were permeabilized with 0.1% Triton X-100 in PBS at room temperature for 15 minutes, followed by blocking with PBS containing 2% FBS and 1% BSA for 2 hours at room temperature. Primary antibodies were applied overnight at 4°C, and sections were then exposed to secondary antibodies at room temperature for 2 hours.

Secondary antibodies used were donkey highly cross-absorbed H + L antibodies conjugated to Alexa 488 and Alexa 568 diluted at 1:2000. Nuclei were counterstained with 0.1 μg/ml DAPI (Sigma). Finally, stained tissues were mounted with Fluoromount G and covered with a glass coverslip. All antibody dilutions for tissue staining contained 1% DMSO to facilitate antibody penetration.

### Microscope image acquisition

All images were obtained using a Zeiss LSM 900 confocal microscope (Axio Observer Z1/7) coupled with an Axiocam 705 camera and immersion objective (Plan-Apochromat 63x/1.4 Oil DIC M27). The images were acquired using ZEN 3.4 (blue edition) software, and visualizations and analyses were performed using ImageJ Fiji.

### Research standards for animal studies

Mice were maintained under specific pathogen-free conditions at the University of Dundee (UK). All animal experiments were ethically reviewed and conducted in compliance with the Animals (Scientific Procedures) Act 1986 and guidelines established by the University of Dundee and the U.K. Home Office. Ethical approval for animal studies and breeding was obtained from the University of Dundee ethical committee, and all procedures were performed under a U.K. Home Office project license. The mice were group-housed in an environment with controlled ambient temperature (20–24°C) and humidity (45–55%), following a 12-hour light/12-hour dark cycle, with ad libitum access to food and water.

## Author contribution (in alphabetical order)

Conceptualization: CYC, DRA, ES, SRP

Investigation: AA, AZ, CF, CYC, NP, WMY, YL

Materials: FT, PLS

Data analysis: AZ, CB, CF, CK, CYC, DWD, IR, JS, MR, NP, OAR, OP, PB, TL, YL, ZKW

Visualization: CYC, NP, SRP

Supervision: DRA, ES, SRP

Writing—original draft: CYC, SRP

Writing—review & editing: AZ, CYC, DRA, ES, SRP

## Data and materials availability

All reagents used in this study are available from commercial sources or repositories without restriction and RRIDs for all reagents are provided in the Key Resource Table. All primary data have been deposited in Zenodo and can be found at https://doi.org/10.5281/zenodo.14911979.

## Supplemental Materials

**Supplemental Figure 1.**
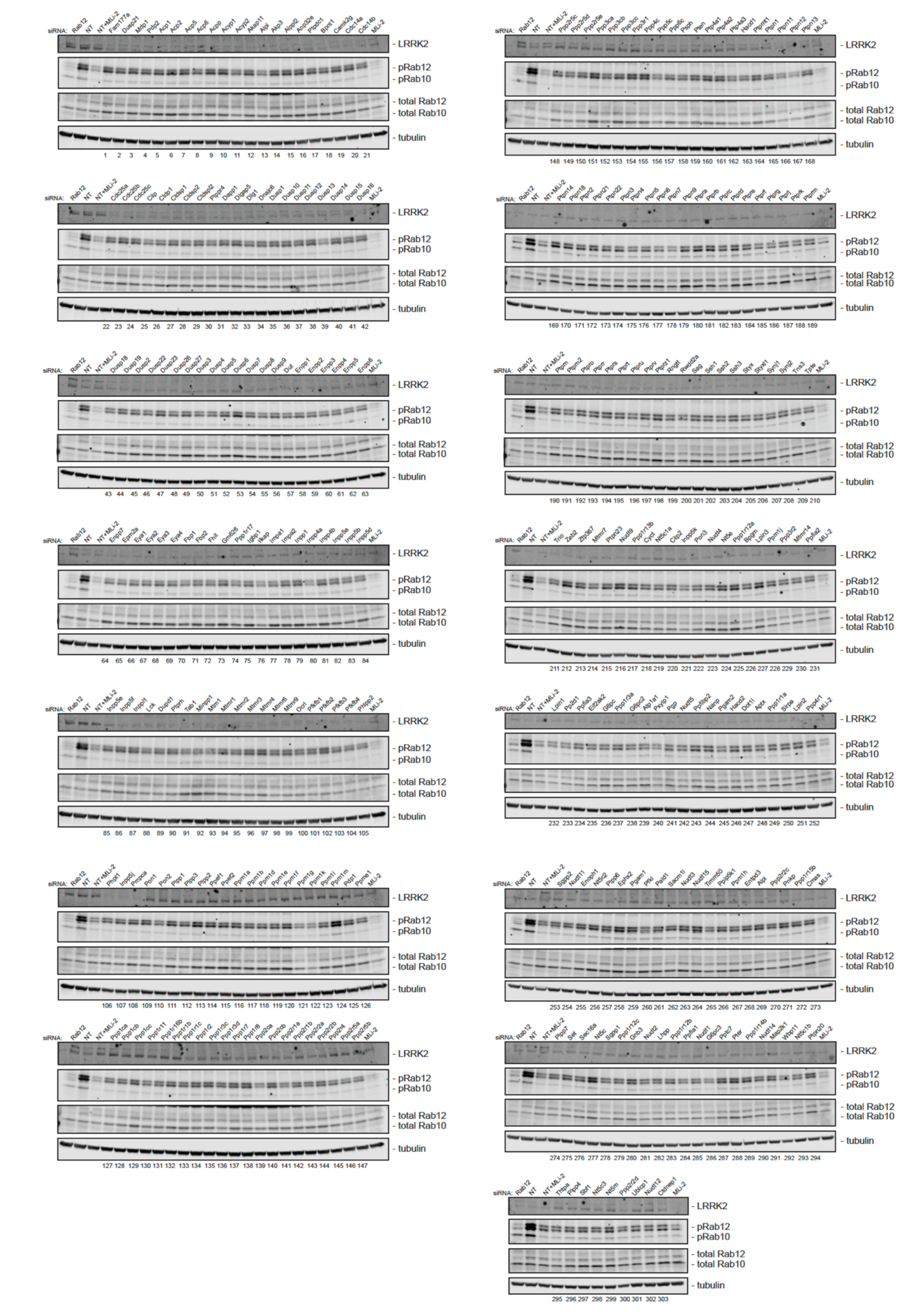
Immunoblots used for screen quantitation. Each lane represents a unique siRNA from the library or Rab12 siRNA, non-targeting siRNA, or non-targeting siRNA with MLi-2. siRNAs were identified by ID number only (labeled below each blot) during screen and analysis. A list of all genes and guides is included in Supplemental Table 2.

**Supplemental Figure 2.**
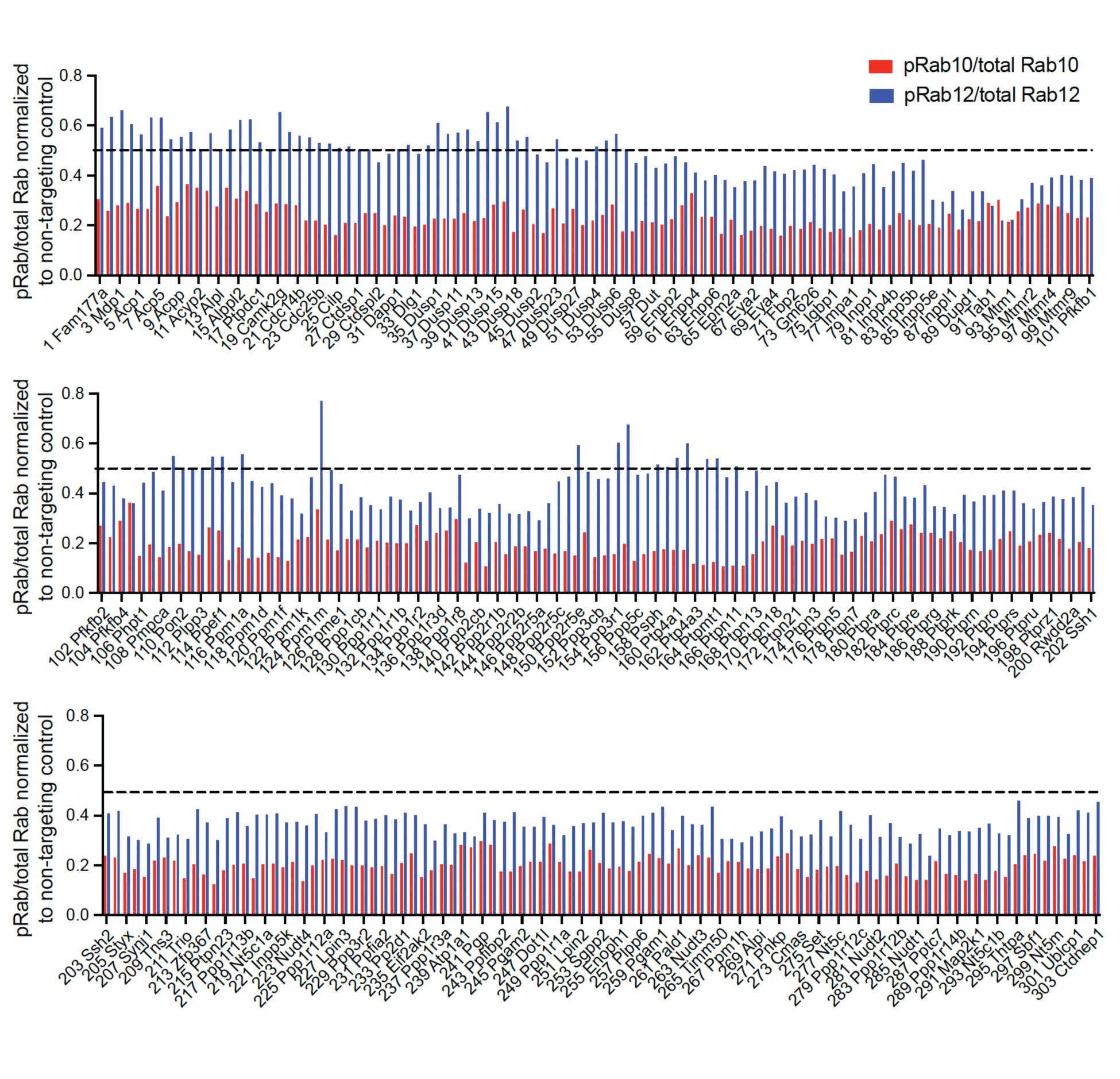
Quantitative analysis of the phosphatome-wide screen. Quantitation of pRab12/total Rab12 and pRab10/total Rab10 from gels in Supplemental Figure 1, normalized to non-targeting (NT) control. Values are the average of two independent analyses of the same data.

**Supplemental Figure 3.**
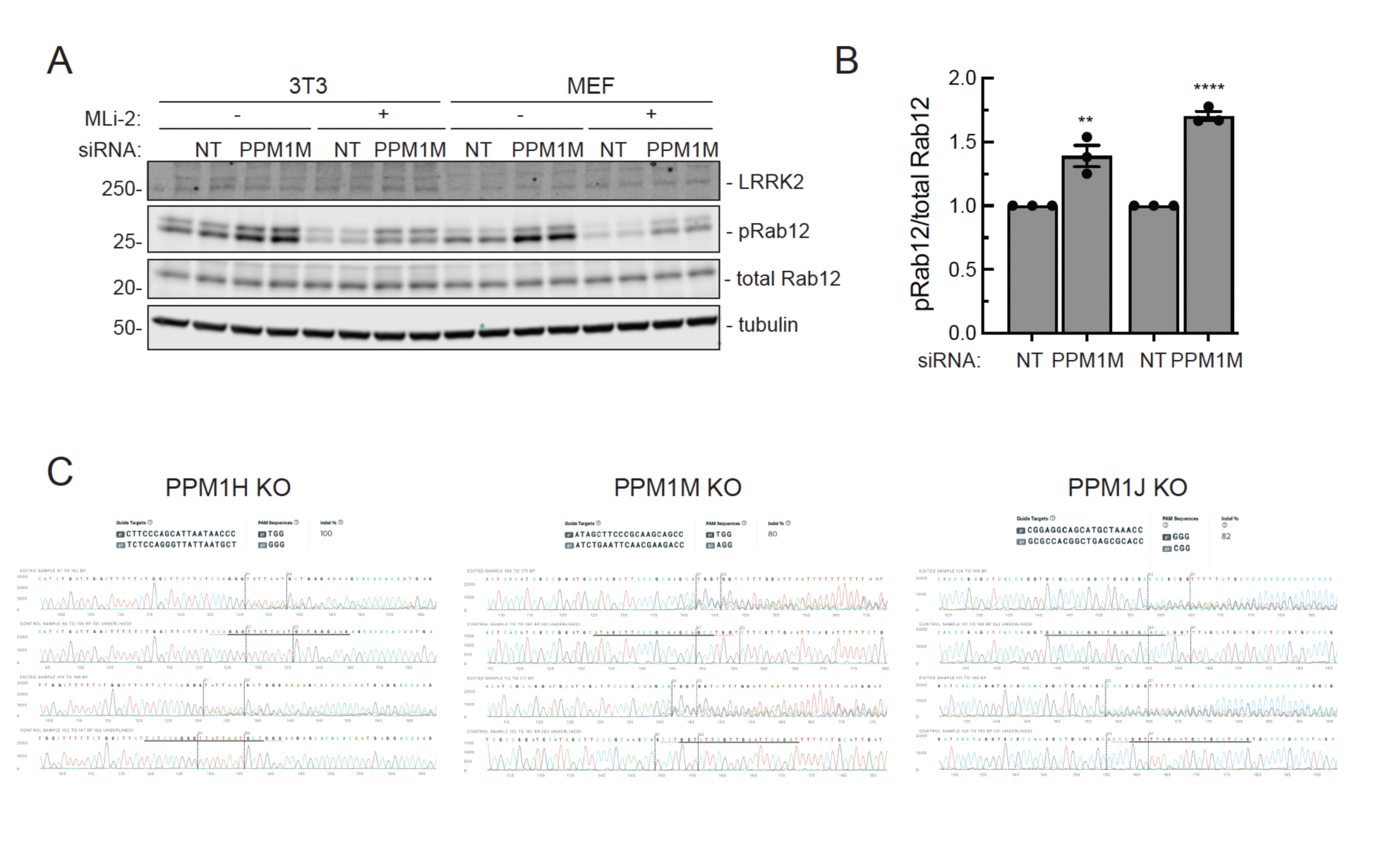
PPM1M knockdown in 3T3 and MEF cells increases pRab12 levels. **(A)** Immunoblot analysis of 3T3 and MEF cells treated with non-targeting (NT) or PPM1M siRNA for 72h, followed by 200 nM MLi-2 for 20 minutes as indicated. **(B)** Quantitation of pRab12 levels from immunoblots in (A) normalized to respective NT controls. Error bars indicate SEM from three independent experiments carried out in duplicate. Statistical significance determined by student’s T-test, respective to NT. **p=0.0093 for 3T3, ****p<0.0001 for MEF. **(C)** Genotyping results of pooled CRISPR knockouts for *Ppm1h*, *Ppm1m*, and *Ppm1j* MEFs by Synthego ICE software. For each cell line, guide sequences, indel frequency (%), and sequencing traces for edited and control samples for each guide are shown.

**Supplemental Figure 4.**
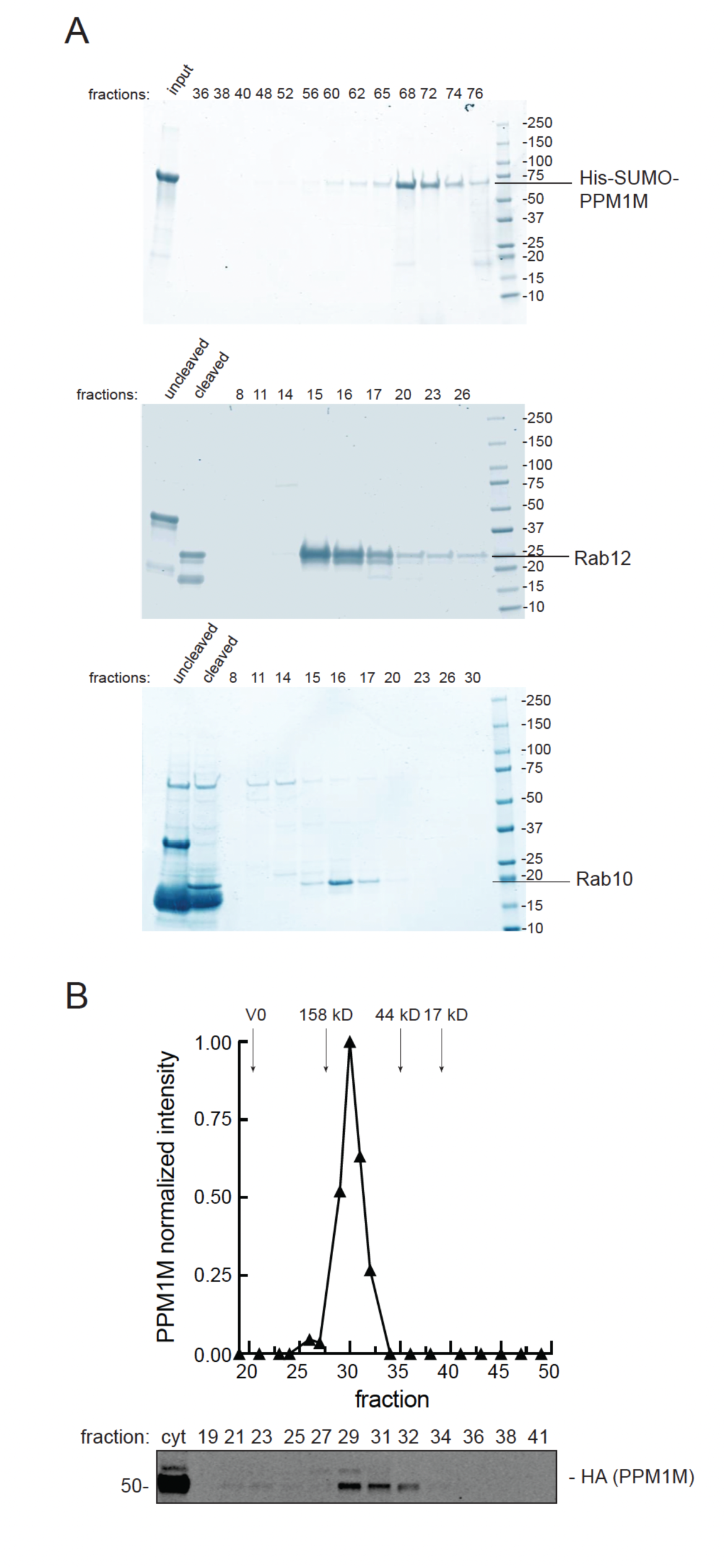
**(A)** SDS-PAGE elution profiles for His-SUMO-PPM1M after purification on a Superdex 200 16/60 120 mL column (top row) and SUMO-tag cleaved (His-SUMO-)Rab10 (middle panel) or (His-SUMO-)Rab12 (bottom panel) after purification on a Superdex 200 10/300 24 mL column. Fraction 72 was used for His-SUMO-PPM1M in experiments. Mass is shown at right in kDa. **(B)** PPM1M chromatographs as a dimer in cytosol. HEK293 cytosol overexpressing HA-PPM1M was resolved on a Superdex 200 10/300 24 mL FPLC column. Immunoblot of fractions collected and their quantitation is shown.

**Supplemental Table 1.** Key Resource Table.

| RESOURCE TYPE | RESOURCE NAME | SOURCE | IDENTIFIER | NEW/ REUSE | ADDITIONAL INFORMATION |
| --- | --- | --- | --- | --- | --- |
| Antibody | anti-GAPDH<br>(mouse monoclonal) | Santa Cruz Biotechnology | sc-32233<br>(RRID: AB_627679) | REUSE | (1:5000) |
| Antibody | anti-PPM1H<br>(rabbit monoclonal) | Abcam | ab303536<br>(RRID: AB_2941812) | REUSE | (1:2000) |
| Antibody | anti-alpha tubulin<br>(mouse monoclonal) | Santa Cruz Biotechnology | sc-32293<br>(RRID: AB_628412) | REUSE | (1:4000) |
| Antibody | anti-HA<br>(rabbit polyclonal) | Sigma | H6908<br>(RRID: AB_260070) | REUSE | (1:1000) |
| Antibody | anti-HA<br>(mouse monoclonal) | Sigma | H9658<br>(RRID: AB_260092) | REUSE | (1:1000) |
| Antibody | anti-HA<br>(rat monoclonal) | Sigma | 11867423001<br>(RRID: AB_390918) | REUSE | (1:1000) |
| Antibody | anti-LRRK2<br>(mouse monoclonal) | Antibodies Incorporated/N euroMab | N241A/34<br>(RRID: AB_10675136) | REUSE | (1:1000) |
| Antibody | anti-LRRK2 pS935<br>(rabbit monoclonal) | MRC PPU U. Dundee and/or Abcam Inc. | UDD2/<br>ab133450<br>(RRID: AB_2732035) | REUSE | (1:1000) |
| Antibody | anti-Rab10 pThr73<br>(rabbit monoclonal) | Abcam Inc. | ab230261<br>(RRID: AB_2811274) | REUSE | (1:1000) |
| Antibody | anti-Rab10<br>(mouse monoclonal) | Abcam Inc. | ab104859<br>(RRID: AB_10711207) | REUSE | (1:1000) |
| Antibody | anti-Rab12 pSer106<br>(rabbit monoclonal) | Abcam Inc. | ab256487<br>(RRID: AB_2884880) | REUSE | (1:1000) |
| Antibody | anti-Rab12<br>(mouse monoclonal) | Santa Cruz Biotechnology | sc-515613<br>(RRID: AB_3101762) | REUSE | (1:500) |
| Antibody | IRDye 800CW Donkey anti-Rabbit IgG | LI-COR | 926-32213<br>(RRID: AB_621848) | REUSE | (1:10,000) |
| Antibody | IRDye 680RD Donkey anti-Rabbit IgG | LI-COR | 926-68073<br>(RRID: AB_10954442) | REUSE | (1:10,000) |
| Antibody | IRDye 800CW Donkey anti-Mouse IgG | LI-COR | 926-32212<br>(RRID: AB_621847) | REUSE | (1:10,000) |
| Antibody | IRDye 680RD Donkey anti-Mouse IgG | LI-COR | 926-68072 | REUSE | (1:10,000) |
|  |  |  | (RRID:<br>AB_10953628) |  |  |
| Antibody | anti-Choline<br>Acetyltransferase<br>(goat polyclonal) | Sigma | 208371<br>(RRID:AB_2079751) | REUSE | (1:200) |
| Antibody | anti-Adenylate cyclase<br>III<br>(rabbit polyclonal) | EnCOR | RPCA-ACIII<br>(RRID:AB_2572219) | REUSE | (1:10,000) |
| Antibody | H+L Donkey anti-Goat<br>AF 488 | Life<br>Technologies | A11055<br>(RRID:AB_2534102) | REUSE | (1:2000) |
| Antibody | H+L Donkey anti-Rabbit AF 568 | Life<br>Technologies | A10042<br>(RRID:AB_2534017) | REUSE | (1:2000) |
| Bacterial strain | MAX Efficiency™<br>DH5α Competent Cells | Invitrogen | 18258012 | REUSE |  |
| Bacterial strain | BL21(DE3)pLysS | Novagen | 69451-3 | REUSE |  |
| Chemical,<br>peptide, or<br>recombinant<br>protein | MLi-2 | MRC PPU<br>Reagents and<br>Services,<br>U. Dundee | CAS No.: 1627091-47-7 | REUSE |  |
| Chemical,<br>peptide, or<br>recombinant<br>protein | Instant Blue<br>Coomassie | Abcam | ab119211 | REUSE |  |
| Chemical,<br>peptide, or<br>recombinant<br>protein | Bio-Rad Protein Assay<br>Dye Reagent<br>Concentrate | Bio-Rad | 5000006 | REUSE |  |
| Chemical,<br>peptide, or<br>recombinant<br>protein | Dharmafect 1 | Dharmacon | T-2001 | REUSE |  |
| Chemical,<br>peptide, or<br>recombinant<br>protein | DMEM (high glucose) | Cytiva | SH30243.02 | REUSE |  |
| Chemical,<br>peptide, or<br>recombinant<br>protein | Fetal bovine serum | Sigma | F0926 | REUSE |  |
| Chemical, peptide, or recombinant protein | Penicillin-Streptomycin | Sigma | P4333 | REUSE |  |
| Chemical, peptide, or recombinant protein | Opti-MEM | Gibco | 31985088 | REUSE |  |
| Chemical, peptide, or recombinant protein | PEI, 25 kDa | Polysciences | 23966 | REUSE |  |
| Chemical, peptide, or recombinant protein | cOmplete EDTA-free protease inhibitor cocktail | Roche | 11873580001 | REUSE |  |
| Chemical, peptide, or recombinant protein | PhosSTOP phosphatase inhibitor cocktail | Roche | 4906837001 | REUSE |  |
| Chemical, peptide, or recombinant protein | Microcystin-LR | Sigma | 475815-M | REUSE |  |
| Chemical, peptide, or recombinant protein | Mouse Rab12 siRNA (OnTarget, SMARTpool) | Dharmacon | L-040865-01 | REUSE |  |
| Chemical, peptide, or recombinant protein | Non-targeting siRNA (OnTarget, SMARTpool) | Dharmacon | D-001810-10 | REUSE |  |
| Chemical, peptide, or recombinant protein | Mouse phosphatase siRNA library (OnTarget, SMARTpool) | Dharmacon | G-113705, see Supplemental Table 2 for gene list | REUSE |  |
| Chemical, peptide, or recombinant protein | Cherry-picked mouse phosphatase siRNA library (OnTarget, SMARTpool) | Dharmacon | custom, see Supplemental Table 2 for gene list | NEW | Additional phosphatase library genes not included in commercial phosphatase library |
| Chemical, peptide, or | GoTaq Green Master Mix | Promega | M7122 | REUSE |  |
| recombinant protein |  |  |  |  |  |
| Critical commercial assay | 4–20% Criterion TGX Precast Midi Protein Gel, 26 well, 15 µl | Bio-Rad | 5671095 | REUSE |  |
| Critical commercial assay | Trans-Blot Turbo RTA Midi 0.2 µm Nitrocellulose Transfer Kit | Bio-Rad | 1704271 | REUSE |  |
| Critical commercial assay | HiTrap TALON crude 1 mL column | Cytiva | 28953766 | REUSE |  |
| Critical commercial assay | Econospin column | Epoch Lifesciences | 1920-050/250 | REUSE |  |
| Critical commercial assay | Electroporation Cuvettes, 0.2 cm gap | Bio-Rad | 1652086 | REUSE |  |
| Dataset | PPM1M, a LRRK2-counteracting, phosphoRab12-preferring phosphatase with potential link to Parkinson's disease | Zenodo | https://doi.org/10.5281/zenodo.14911979 | NEW | raw tabular data and original tiff files for all western blot analysis, raw tabular data and original microscopy images for ciliation experiments |
| Experimental model: Cell line | HEK293T (human) | ATCC | CRL-3216 (RRID: CVCL_0063) | REUSE |  |
| Experimental model: Cell line | A549 (human) | ATCC | CCL-185 (RRID: CVCL_0023) | REUSE |  |
| Experimental model: Cell line | PPM1H knockout A549 (human) | MRC PPU Reagents and Services, U. Dundee | PMIID: 31663853 | REUSE |  |
| Experimental model: Cell line | Mouse Embryonic Fibroblasts (mouse) | MRC PPU Reagents and Services, U. Dundee | RRID: CVCL_E7DI | NEW | littermate match to PPM1M knockout MEFs |
| Experimental model: Cell line | PPM1M knockout Mouse Embryonic Fibroblasts (mouse) | MRC PPU Reagents and Services, U. Dundee | RRID: CVCL_E7DI | NEW | littermate match to WT MEFs |
| Experimental model: Cell line | 3T3 Flp In (mouse) | Invitrogen | R76107<br>(RRID: CVCL_U422) | REUSE |  |
| Protocol | Gibson Assembly | [protocols.io](https://doi.org/10.17504/protocols.io.eq2lyjwyqlx9/v1) | https://doi.org/10.17504/protocols.io.eq2lyjwyqlx9/v1 | NEW |  |
| Protocol | Phosphatome-wide siRNA screen in 3T3 cells | [protocols.io](https://doi.org/10.17504/protocols.io.36wgqdxr5vk5/v1) | [dx.doi.org/10.17504/protocols.io.36wgqdxr5vk5/v1](https://doi.org/10.17504/protocols.io.36wgqdxr5vk5/v1) | NEW |  |
| Protocol | Creation of pooled CRISPR KO cell lines using Synthego sgRNA | [protocols.io](https://doi.org/10.17504/protocols.io.bp2l6dqodvqe/v1) | [dx.doi.org/10.17504/protocols.io.bp2l6dqodvqe/v1](https://doi.org/10.17504/protocols.io.bp2l6dqodvqe/v1) | REUSE |  |
| Protocol | Isolation of mouse embryonic fibroblasts (MEFs) from mouse embryos at E12.5 | [protocols.io](https://doi.org/10.17504/protocols.io.eq2ly713qlx9/v1) | [dx.doi.org/10.17504/protocols.io.eq2ly713qlx9/v1](https://doi.org/10.17504/protocols.io.eq2ly713qlx9/v1) | REUSE |  |
| Protocol | Quantitative Immunoblotting Analysis of LRRK2 Signalling Pathway | [protocols.io](https://doi.org/10.17504/protocols.io.bsgnrbv6) | [dx.doi.org/10.17504/protocols.io.bsgnrbv6](https://doi.org/10.17504/protocols.io.bsgnrbv6) | REUSE |  |
| Protocol | Expression and purification of PPM1H phosphatase | [protocols.io](https://doi.org/10.17504/protocols.io.bu7wnzpe) | [dx.doi.org/10.17504/protocols.io.bu7wnzpe](https://doi.org/10.17504/protocols.io.bu7wnzpe) | REUSE |  |
| Protocol | Dephosphorylation of phosphorylated Rab GTPases by PPM phosphatases | [protocols.io](https://doi.org/10.17504/protocols.io.5jyl8d4j7g2w/v1) | [dx.doi.org/10.17504/protocols.io.5jyl8d4j7g2w/v1](https://doi.org/10.17504/protocols.io.5jyl8d4j7g2w/v1) | NEW |  |
| Protocol | Crude Membrane Fractionation of Cultured Cells | [protocols.io](https://doi.org/10.17504/protocols.io.yxmvmnb99g3p/v1) | https://doi.org/10.17504/protocols.io.yxmvmnb99g3p/v1 | REUSE |  |
| Protocol | Expression and purification of Rab8A (1-181) stoichiometrically phosphorylated at pThr72 | [protocols.io](https://doi.org/10.17504/protocols.io.butinwke) | [dx.doi.org/10.17504/protocols.io.butinwke](https://doi.org/10.17504/protocols.io.butinwke) | REUSE |  |
| Protocol | Analysis of Primary Cilia in Rodent Brain By Immunofluorescence Microscopy | [protocols.io](https://doi.org/10.17504/protocols.io.bnwmfc) | [dx.doi.org/10.17504/protocols.io.bnwmfc](https://doi.org/10.17504/protocols.io.bnwmfc) | REUSE |  |
| Recombinant DNA | pCMV5D HA-PPM1H | MRC PPU Reagents and Services, | DU62789 | REUSE |  |
|  |  | U. Dundee |  |  |  |
| Recombinant DNA | pCMV5D HA-PPM1H H153D | MRC PPU Reagents and Services, U. Dundee | DU62928 | REUSE |  |
| Recombinant DNA | pCMV5D HA-PPM1H D288A | MRC PPU Reagents and Services, U. Dundee | DU62985 | REUSE |  |
| Recombinant DNA | pCMV5D HA-PPM1M | MRC PPU Reagents and Services, U. Dundee | DU68124 | REUSE |  |
| Recombinant DNA | pCMV5D HA-PPM1M H127D | MRC PPU Reagents and Services, U. Dundee | DU68165 | REUSE |  |
| Recombinant DNA | pCMV5D HA-PPM1M D235A | MRC PPU Reagents and Services, U. Dundee | DU68164 | REUSE |  |
| Recombinant DNA | pCMV5D HA-PPM1M D440N | MRC PPU Reagents and Services, U. Dundee | DU72159 | NEW |  |
| Recombinant DNA | pCMV5D HA-PPM1H_M flap | Addgene | Addgene in progress | NEW |  |
| Recombinant DNA | pCMV5D HA-PPM1M_H flap | Addgene | Addgene in progress | NEW |  |
| Recombinant DNA | Flag-LRRK2 R1441C | MRC PPU Reagents and Services, U. Dundee | DU13078 | REUSE |  |
| Recombinant DNA | Flag-LRRK2 R1441G | MRC PPU Reagents and Services, U. Dundee | DU26477 | REUSE |  |
| Recombinant DNA | His-SUMO-PPM1M | MRC PPU Reagents and Services, U. Dundee | DU68141 | REUSE |  |
| Recombinant DNA | His-SUMO-PPM1M H127D | MRC PPU Reagents and Services, U. Dundee | DU68200 | REUSE |  |
| Recombinant DNA | His-SUMO-PPM1M D440N | MRC PPU Reagents and Services, U. Dundee | DU72158 | REUSE |  |
| Recombinant DNA | His-SUMO-PPM1H | MRC PPU Reagents and Services, U. Dundee | DU62835 | REUSE |  |
| Recombinant DNA | His-SUMO-PPM1H D288A | MRC PPU Reagents and Services, U. Dundee | DU68087 | REUSE |  |
| Recombinant DNA | His-Thrombin-Rab8A (1-181) | MRC PPU Reagents and Services, U. Dundee | DU68198 | REUSE |  |
| Recombinant DNA | His-SUMO-Rab10 Q68L | Addgene | Addgene in progress | NEW |  |
| Recombinant DNA | His-SUMO-Rab12 Q101L | Addgene | 208371 (RRID:Addgene_208371) | REUSE |  |
| Software/code | AlphaFold Server | AlphaFold 3 | RRID: SCR_025885 | REUSE |  |
| Software/code | ChimeraX | ChimeraX | PMID: 32881101 (RRID: SCR_015872) | REUSE |  |
| Software/code | ImageJ | ImageJ version 2.14 | RRID: SCR_003070 | REUSE |  |
| Software/code | Graphpad Prism | Prism 10 version 10.2.3 | RRID: SCR_002798 | REUSE |  |
| Software/code | ZEN | Zeiss ZEN Microscopy Software | RRID:SCR_013672 | REUSE |  |

**Supplemental Table 2. List of siRNA targets and sequences of all siRNA oligonucleotides used in phosphatome-wide siRNA screen.** 4 siRNAs per gene were provided in SMARTpool format. Of note, the Dharmacon commercial mouse phosphatase library did not include all phosphatase genes; we also created a custom library of 89 additional phosphatase genes. These are described respectively as G-113705 Lot 23107 (commercial library) or “cherry pick” library in the table. During siRNA screen and analysis, samples were identified only by the gene ID numbers listed.

**Supplemental Table 3.** Overview of cohorts that were interrogated for *PPM1M* p.D440N carrier status. Included is the respective cohort including total number of individuals per study, number of disease and control subjects, number of individuals carrying the *PPM1M* D440N variant in the heterozygous state with and without PD, and minor allele frequency (MAF). YOPD = young onset PD.

| Study | N | Disease / control cohort (if applicable) | *PPM1M* D440N PD | *PPM1M* D440N non-PD | MAF |
| --- | --- | --- | --- | --- | --- |
| Hop et. al | 71959 | 2184 (familial PD cases) / 69775 controls | 3 | 3 | 3.81E-05 |
| Austrian PD cohort | 382 | 382 (familial and YOPD cases) | 1 | n.a. | - |
| Mayo clinic | 700 | 700 (neuropathological specimens) | 1 | n.a. | - |
| Czechia PD cases | 33 | 33 | 0 | n.a. | - |
| Ireland PD cases | 672 | 672 | 0 | n.a. | - |
| Poland PD cases | 725 | 725 | 1 | n.a. | - |
| Ukraine PD cases | 139 | 139 | 1 | n.a. | - |
| gnomAD 4.1 | 800000 | 800000 (general population) | N/A | 66 | 4.13E-05 |
| Centogene | 192000 | 10000 PD cases / 182000 non-PD cases | 0 | 13 | 3.57E-05 |

